# Influenza virus antagonizes self sensing by RIG-I to enhance viral replication

**DOI:** 10.1101/2025.03.12.642847

**Authors:** Mitchell P. Ledwith, Thomas Nipper, Kaitlin A. Davis, Deniz Uresin, Zhenyu Zhang, Mariel Kleer, Denys A. Khaperskyy, Craig McCormick, Anastassia V. Komarova, Andrew Mehle

**Affiliations:** Medical Microbiology and Immunology, University of Wisconsin Madison, Madison, WI, USA; Institut Pasteur, Université Paris Cité, Interactomics, RNA and Immunity laboratory, F-75015 Paris, France; Department of Microbiology & Immunology, Dalhousie University, Halifax, Nova Scotia, Canada; Institut Pasteur, Université Paris Cité, Molecular Genetics of RNA Viruses, CNRS UMR-3569, F-75015 Paris, France; Institut Pasteur, Pasteur-Oncovita Joint Laboratory, F-75015 Paris, France

**Keywords:** innate immunity, self sensing, RIG-I, RNA polymerase III, influenza virus

## Abstract

Innate immune sensors must finely distinguish pathogens from the host to mount a response only during infection. RIG-I is cytoplasmic sensor that surveils for foreign RNAs. When activated, RIG-I triggers a broad antiviral response that is a major regulator of RNA virus infection. Here were show that RIG-I not only bound viral RNAs, but was activated by host RNAs to amplify the antiviral state. These were primarily non-coding RNAs transcribed by RNA polymerase III. They were benign under normal conditions but became immunogenic during influenza virus infection where they signaled via RIG-I to suppress viral replication. This same class of RNAs was bound by influenza virus nucleoprotein (NP), which normally functions to encapsidate the viral genome. NP interacted with RIG-I and cytoplasmic NP antagonized sensing of self RNAs to counter innate immune responses. Overall, these results demonstrate that self sensing is strategically deployed by the cell to amplify the antiviral response and reveal a newly identified viral countermeasure that disrupts RIG-I activation by host RNAs.

## Introduction

Cells employ complex sensing pathways to detect nucleic acids from invading pathogens and initiate innate immune responses^1–3^. Foreign nucleic acids exhibit unique molecular features and subcellular localizations that allow nucleic acid sensors to differentiate them from host nucleic acids. Nucleic acid sensors use these features to carefully balance sensitivity and specificity for rapid and robust responses only during infection. Whereas foreign nucleic acid sensing is protective, inappropriate sensing of self nucleic acids can cause chronic inflammation and immunopathology^4^.

RIG-I-like receptors (RLRs) are germline encoded RNA sensors composed of RIG-I (*DDX58*), MDA5 (*IFIH1*) and LGP2 (*DHX58*) that surveil the cytoplasm for foreign RNAs^1,2^. MDA5 recognizes long dsRNAs (>0.5kb), a common intermediate during RNA virus genome replication and a by-product of bi-directional transcription for certain DNA viruses^5^. RIG-I recognizes shorter dsRNA structures and requires the presence of a 5’ diphosphate (5’ PP) or 5’ triphosphate (5’ PPP) for activation^6–8^. The 5’ PP/PPP moieties are products of replication by viral RNA-dependent RNA polymerases. When activated, both RIG-I and MDA5 undergo conformational changes that expose their tandem caspase recruitment domains (CARD)^9–12^. The CARDs then oligomerize and signal through MAVS to initiate an anti-pathogen response by expressing type I and type III interferons (IFN) and other proinflammatory cytokines.

RIG-I is constitutively expressed to enable detection of the earliest stages of viral infection, including sensing of viral genomes even before their replication or transcription has begun^13,14^. RIG-I is the dominant innate immune sensor controlling influenza virus replication where it binds and is activated by viral genomes and aberrant replication products^15–19^. However, RIG-I does not exclusively bind to viral RNAs; it samples dsRNAs regardless of their immunogenicity or origin^20,21^. The total numbers of RNA molecules under surveillance vastly outnumbers the amount of RIG-I, and only a fraction of those are viral RNAs. RIG-I must therefore rapidly recycle off non-immunogenic RNAs so it can continue sampling all available RNAs to detect the rare immunogenic species. Discrimination occurs after RIG-I binds dsRNA where it has a faster off rate and lower affinity for dsRNA lacking a 5’ PPP that promotes translocation along the length of the RNA and rapid recycling^12,22–24^. Conversely, when RIG-I encounters dsRNA with a 5’ PP/PPP, it assumes a distinct conformation with high-affinity interactions between the RNA and the 5’ PPP^12^. This throttles RIG-I translocation helping to form long-lived complex that promote RIG-I oligomerization and maintain the CARD domains in an exposed state competent to form a signaling platform with MAVS^9,11,12,22^. These subtle differences in RNA binding and occupancy time establish mechanisms that distinguish virus from host while also allowing limiting amounts of RIG-I to surveil vast cytoplasmic RNAs.

Aberrant sensing of self RNAs caused by inborn genetic errors in RIG-I or MDA5 is associated with multiple disease states including interferonopathies like Singleton-Merten syndrome and Aicardi–Goutières syndrome^4^. This failure to faithfully differentiate self from non-self leads to chronic interferon signaling and associated inflammation. While self sensing has been generally viewed as pathogenic, new data suggest that it may also amplify antiviral responses^25–29^. To define a potential protective role of self sensing during viral infection, we quantitatively characterized RNAs surveilled by RIG-I under sterile conditions and during viral infection. The vast majority of RNAs bound by RIG-I are host RNAs, even during infection. RIG-I-bound host RNAs are dominated by small non-coding RNA polymerase III (Pol III) transcripts, including Y RNAs, vault RNAs (vtRNAs), and U6 RNAs. These RNAs are benign in uninfected cells but become immunogenic during influenza virus infection when they signal via RIG-I to suppress viral replication. This virus:host conflict creates an environment ripe for the emergence of viral countermeasures. We show that influenza virus nucleoprotein (NP), a viral protein whose primary function is to encapsidate the viral genome, binds many of the same host RNAs as RIG-I. NP interacts with RIG-I and antagonizes sensing of self RNAs to counter innate immune responses. RIG-I activation is finely balanced to promote rapid detection of invading pathogens while avoiding self sensing that can cause diseases. Our results show that this balance is intentionally disrupted by cells where RIG-I is activated by both host and viral RNAs to provide parallel mechanisms of innate immune activation during influenza virus infection. This suggests a generalizable mechanism by which the nucleoproteins of RNA viruses could suppress infection-induced immunogenicity triggered by host non-coding RNAs.

## Results

### RIG-I primarily engages host RNAs even during infection

Understanding how the repertoire of RIG-I-bound RNAs changes requires quantitative comparisons across conditions. To achieve this, we developed a spike-in cross-linking immunoprecipitation sequencing (CLIP-Seq) approach. First, accurate quantification and comparisons between immunoprecipitated libraries necessitate equivalent amounts of target protein in each sample, yet this is complicated by the fact that RIG-I expression increases in response to infection^1^. We eliminated this unwanted variable by generating human A549 lung cells that stably express similar amounts of RIG-I in all conditions (Fig 1A). Inoculation of these cells with influenza A virus (FLUAV) or vesicular stomatitis virus (VSV) demonstrated that constitutive RIG-I expression restricts replication of both of these RNA viruses (Fig 1A), as expected^19^. These cells were used to generate lysates for CLIP-seq to which we then introduced a spike-in control to mitigate biases present in global analyses of immunoprecipitated samples^30^. Spike-in controls were generated by cross-linking RIG-I to RNA in *Drosophila* S2 cells (Fig S1A). Sequence divergence between *Drosophila* and humans permitted unique identification and quantification of RNAs derived from the common spike-in control (Fig S1B). This strategy disambiguated >99.5% of sequencing reads (Fig S1C), allowing for internal normalization and differential assessment of RNA-binding across samples.

**Figure 1.**
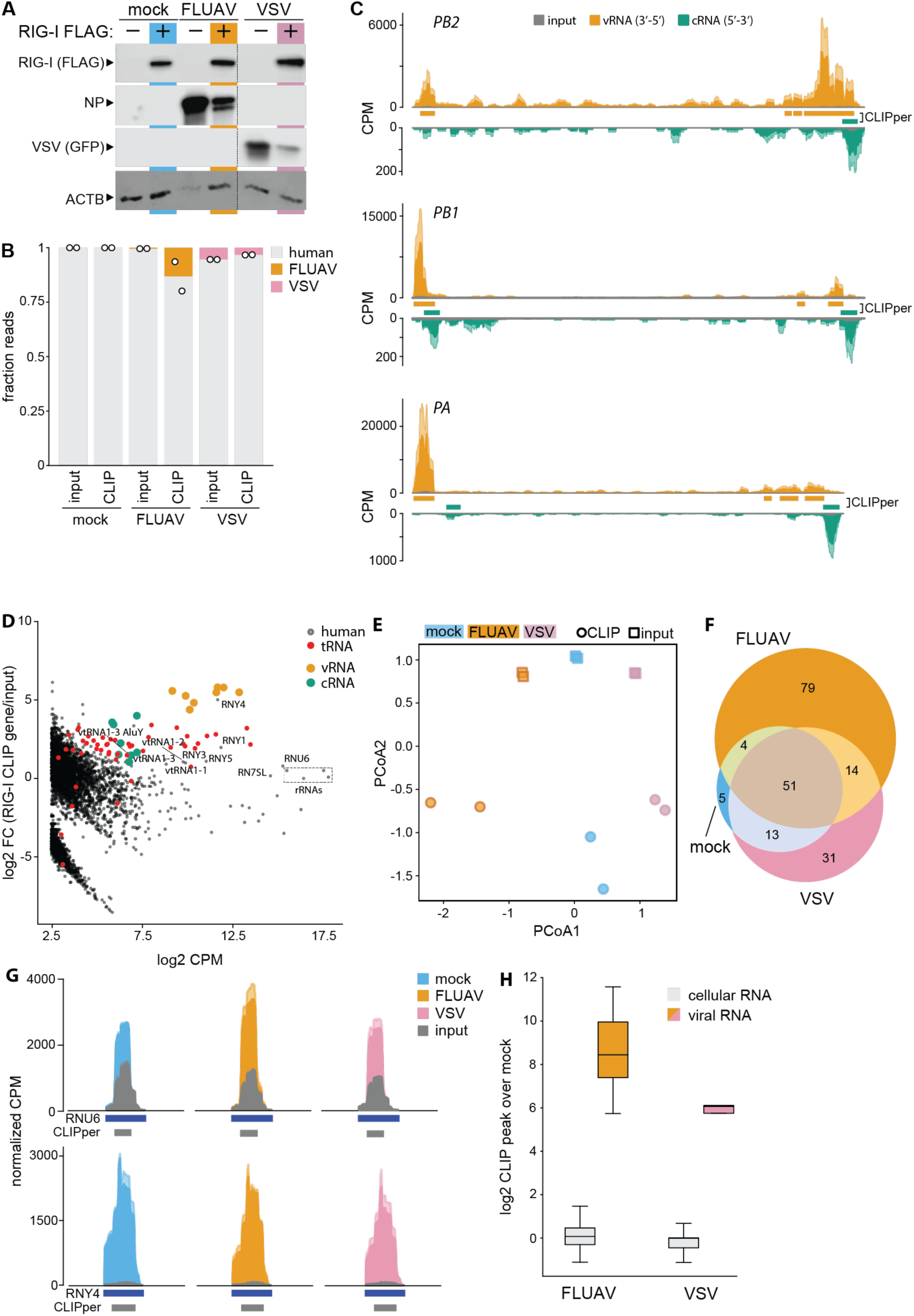
RIG-I primarily surveils host RNAs even during infection. (A) WT A549 cells or those stably expressing RIG-I were mock-infected, infected with FLUAV (MOI = 0.01, 24h), or infected with VSV-GFP (MOI = 0.001, 24h). Protein expression was detected by western blotting. (B) RIG-I CLIP was performed with a spike-in control in biological duplicate. Fraction of reads mapping to human, FLUAV, and VSV genomes for size-matched input controls or RIG-I CLIP samples. (C) Normalized read coverage for RIG-I-bound and size-matched input control RNAs. Data from independent biological replicates are shown as mean and s.d. using dark and light shades of the same color, respectively. CLIPper-identified peaks are shown. CPM, counts per million. See Supplementary Table 1 for all identified RIG-I CLIP peaks. (D) Gene enrichment in RIG-I binding during infection. Log2 fold-change in read depth of genes captured by RIG-I CLIP versus the size-matched input control plotted as a function of the average counts per million (CPM) in the CLIP and size-matched input libraries. Differential gene binding data can be found in Supplementary Table 2. See also Supplementary Table 3 for differential peak binding. (E) Principle coordinate analysis of genes identified in the spike-in-normalized RIG-I CLIP and size-matched input libraries. (F) Overlap of CLIPper-called RIG-I binding peaks from mock, FLUAV-infected, and VSV-infected A549 cells. (G) Normalized read coverage of RIG-bound RNAs or size-matched input controls from mock, FLUAV-infected, and VSV-infected cells shown as in (C). CLIPper-identified peaks and gene tracks are shown. (H) Log2 fold-change of infection over mock for RIG-I CLIP peak intensity separated by host- or virus-derived RNAs. Associated with Supplementary Table 3.

Our quantitative CLIP-Seq approach was used to compare RNAs bound by RIG-I during infection FLUAV, VSV, or mock treatment. As expected, RIG-I bound FLUAV RNAs (Fig 1B-D, Table S1-3). RIG-I binding was localized to the termini of viral genomes, with binding peaks present on both the negative-sense genome (vRNA) and the positive-sense replication intermediate (cRNA) (Fig 1C). This recapitulates prior RIG-I:RNA associations and confirms the known stimulatory role of the double-stranded 5’ PPP panhandle structures that are made by viral genome termini, possibly enriched in defective viral genomes, providing confidence in our approach^16–18^. Influenza virus-derived RNAs constituted ∼15% of the total RNA bound by RIG-I, while VSV-derived RNAs were not even enriched relative to their abundance in the input and were less than 5% of the RIG-I bound RNAs (Fig 1B). The vast majority of RIG-I-bound RNAs across all conditions were derived from the host.

RIG-I bound a discrete population of host non-coding RNAs expressed by Pol III. Non-coding RNAs can be encoded by 1000’s of unique degenerate loci, making them notoriously difficult to quantify and accurately map, especially in CLIP-Seq data^31–34^. Furthermore, these regions may contain additional unannotated transcripts, making mapping to the traditional reference insufficient. We addressed this by using two-pass mapping and de novo transcriptome assembly to quantify these RNA populations (Fig. S1B). Y RNAs (*RNY*), vtRNAs (*vtRNA*), U6 RNAs (*RNU6*), Alu RNAs, and tRNAs were all bound by RIG-I during FLUAV infection (Fig 1D and S2A, Table S2). While ribosomal RNA (rRNAs) are the most abundant RNAs in our samples, they are not enriched by RIG-I confirming the specificity of our CLIP-Seq. Similar patterns were observed during VSV infection and mock-treatment, with RNY4 a notable example of a highly enriched host non-coding (Fig S2A). We confirmed these findings using RNA-immunoprecipitations (RIPs). RIG-I immunoprecipitated the same host non-coding RNAs and PB2, but not a control rRNA (Fig S2B). Principle coordinate analysis cleanly separated RIG-I CLIP-Seq samples from size-matched input controls, reinforcing that RIG-I interacts with a restricted pool of RNAs (Fig 1E). Peak calling identified RNAs differentially bound between mock and infected conditions. Nearly all peaks that were significantly bound in mock-infected cells were also identified in cells infected with influenza virus or VSV infection (Fig 1F, Table S1). RIG-I binds distinct populations in FLUAV- or VSV-infected cells (Fig 1E), yet they share a common core as ∼60% of peaks identified in VSV infection were also identified in FLUAV infection. This overlap may be greater as approximately half of the peaks absent from VSV-infected samples belonged to the influenza virus genome. This shared binding pattern is reflected in the read densities across the RIG-I-bound U6 and Y4 RNAs where binding intensity is consistent between mock, FLUAV-, and VSV-infected cells (Fig. 1G). We extended this analysis to all RNAs bound by RIG-I (Fig 1H, Table S3). RIG-I peaks on viral RNAs were dramatically increased, as expected given their absence in mock conditions. However, host RNAs displayed little change in binding during infection with either FLUAV or VSV. Analysis at the gene level yielded similar trends, with changes in RIG-I binding occurring mostly on viral RNAs (Fig S2C, Table S2). The host transcripts that appear to change RIG-I occupancy during infection are IFN-stimulated genes that are normally absent in mock cells (Table S2), thus changes in binding largely reflect the absence of these transcripts in uninfected cells. Together, these data show that RIG-I continually binds and surveils cellular RNAs independent of viral infection, especially those expressed by Pol III, and that even during infection the majority of RNAs sampled by RIG-I are host-derived.

### Influenza virus NP binds host non-coding RNAs

Like all negative-strand RNA viruses, FLUAV and VSV package their genomes into large ribonucleoprotein complexes (RNPs) with the viral RNA-dependent RNA polymerase (RdRP) bound to the genome termini while the viral nucleocapsid (N) or nucleoprotein (NP) oligomerizes along the length of the genome^35^. RNPs then serve as templates for both genome replication and gene transcription. As both FLUAV NP and RIG-I bind to the viral genome, we asked whether they interact. Cells expressing RIG-I were infected with the 2009 pH1N1 strain of FLUAV or mock treated. NP co-precipitated with RIG-I, but not the control (Fig 2A). This interaction required RNA, as treatment with RNase dramatically reduced co-precipitation of NP by RIG-I (Fig 2B and S3). These findings are reinforced by similar interactions with RIG-I derived from FLUAV hosts like duck and pig^36^.

**Figure 2.**
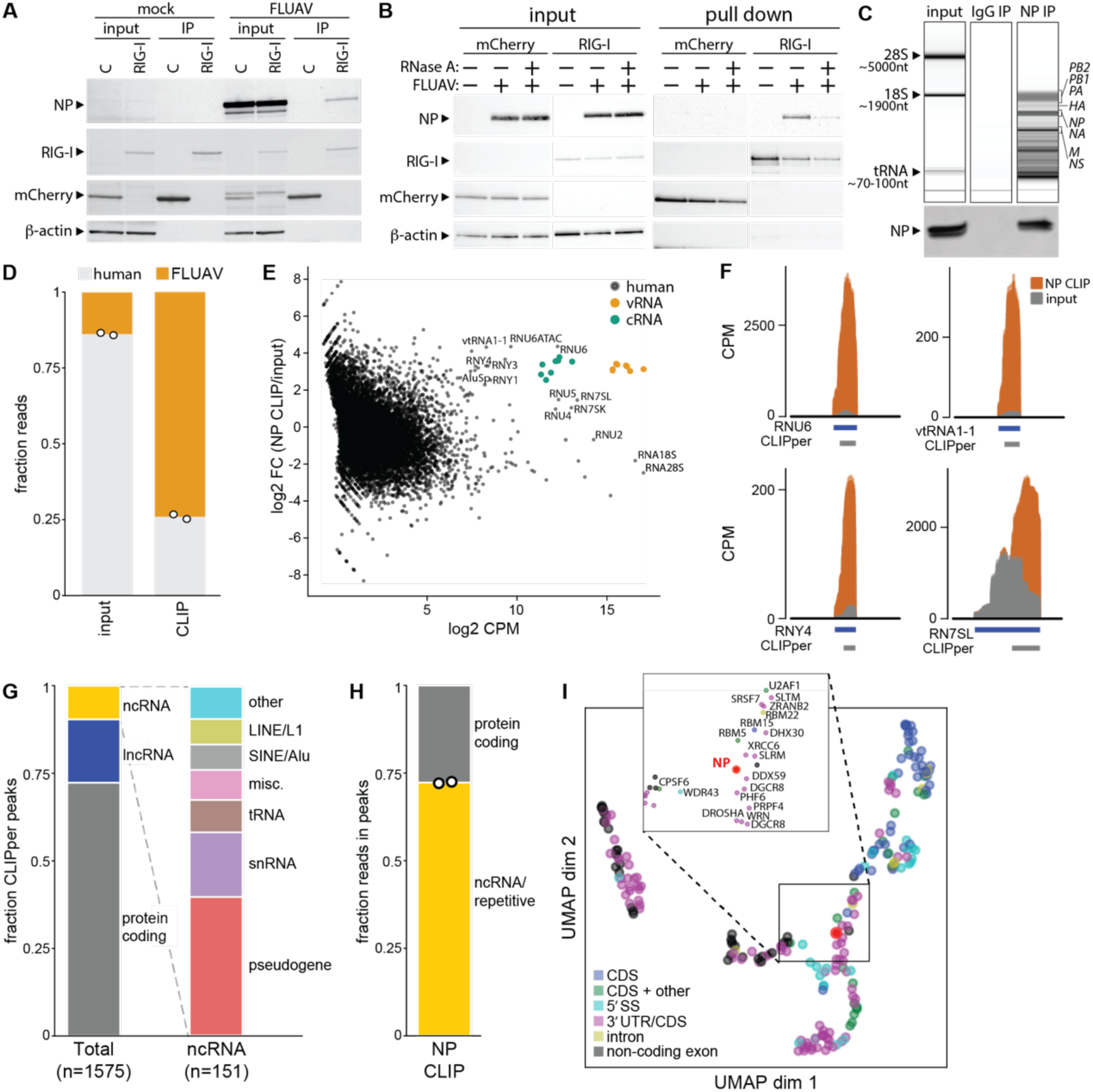
FLUAV NP binds human non-coding RNAs during infection. (A-B) NP interacts with RIG-I in an RNA-dependent manner. Cells expressing Strep-tagged RIG-I or an mCherry control were infected with 2009 pH1N1 influenza virus, lysed and subject to enrichment with Strep-tag affinity resin. Proteins in the input lysate and enriched samples were detected by western blot. In (B), samples were subject to RNAse A treatment prior to affinity capture. (C) RNA-immunoprecipitation was performed on lysates prepared from A549 cells infected for 24h with A/WSN/33 (MOI = 0.02) using antibody recognizing NP or an IgG control. RNAs were subject to Bioanalyzer analysis and NP was detected by western blot. (D) Frequency of host- and FLUAV-derived reads in SMInput and NP eCLIP libraries prepared from A549 cells infected with FLUAV for 24h (MOI = 0.02). (E) Identification of NP-bound gene transcripts during infection. Log2 fold-change in read depth of genes captured by NP CLIP versus the size-matched input control plotted as a function of the average counts per million (CPM) in the CLIP and size-matched input libraries. Negative-sense vRNA and positive-sense cRNA viral gene segments and select host genes are indicated. Differential gene binding data can be found in Supplementary Table 4. (F) Normalized read coverage for NP-bound and size-matched input control RNAs. Data from independent biological replicates are shown as mean and s.d. using dark and light shades of the same color, respectively. CLIPper-identified peaks and gene tracks are shown. CPM, counts per million. See Supplementary Table 5 for all identified NP CLIP peaks. (G) Human RNA biotypes of CLIPper-called peaks (left) and further division of non-coding RNAs (ncRNA) into annotated biotypes (right). (H) Read counts in CLIPper-called peak were assigned to host-derived protein-coding or non-coding/repetitive RNAs based on their RNA biotype. (I) Clustering of eCLIP datasets from ENCORE database with uniform manifold approximation and projection based on Jaccard similarity. Dot color denoted by NP eCLIP library (red) or available hierarchical clustering of eCLIP libraries in the ENCORE database.

NP binds RNA non-specifically, forming sequence-independent interactions with the phosphate backbone allowing it to coat the entirety of the viral genome^37–42^. This raises the possibility that NP interacts with cellular RNAs in addition to the viral genome. NP was immunoprecipitated from infected human A549 lungs cells and sizing analysis of the associated RNA identified species corresponding to viral genomic RNAs (900-2100nt) (Fig 2C). In addition to these larger RNAs, immunoprecipitations also enrich smaller RNA species (100-300nt) that could be aberrant products of genome replication or host-derived RNAs. To profile the full spectrum of RNAs bound by NP in an unbiased fashion, we performed classic CLIP-Seq targeting NP in infected A549 cells. ∼75% of the bound RNAs were derived from the viral genome (Fig 2D). Viral RNAs were dramatically enriched relative to their abundance in the infected cells (Fig 2D-E), confirming our ability to detect known targets of NP. Similar to previous observations, NP bound the viral genome non-uniformly, with specific regions reproducibly bound at higher levels (Fig S4A-B)^43–45^. Surprisingly, ∼25% of bound sequences are host-derived, suggesting that a significant portion of NP engages host RNAs during infection. Gene-level analysis revealed that many of the host RNAs belong to repetitive and non-coding RNA (ncRNA) families, especially vault RNAs, Y RNA, Alu RNAs, the spliceosomal small nuclear RNAs (snRNAs) U5, U6, and U6ATAC, 7SK snRNA, and the 7SL signal recognition particle RNA (Fig 2E-F and S4C, Table S4). To confirm the CLIP results, we performed NP-RIPs and probed for a subset of the host ncRNAs. NP selectively enriched U6, Y4, vtRNA1-1 at levels similar to the PB2 genome segment (Fig S4D). Bound RNAs were frequently Pol III transcripts. 1575 high-confidence binding peaks were identified, of which ∼150 are annotated as non-coding RNAs (Fig 2G). Despite representing only ∼10% of the identified peaks, small non-coding RNAs contributed more than 70% of total reads at NP-bound peaks (Fig 2H). Given the binding preference of NP, these regions are presumably single stranded^41,46^. No obvious sequence motifs were found in NP binding sites on host RNAs, consistent with how NP binds the viral genome in a sequence-independent fashion.

RNA-binding proteins typically have a restricted binding repertoire of one or a few classes of RNAs^33^. Unlike many previously profiled RNA-binding proteins, NP enriched for RNAs spanning several diverse classes and localizing to multiple cellular compartments. To contextualize this unique RNA-binding activity, we compared NP-bound peaks to the publicly available ENCORE database of CLIP datasets^33^. Dimensionality reduction and visualization recapitulated known binding preferences of proteins in the ENCORE database (Fig 2I). NP clustered with RNA-binding proteins with disparate roles, consistent with its diverse RNA-binding profile. Proteins in the NP-proximal embedding space act in the processing of non-coding RNAs, such as DCGR8, RMB5, and WDR43, which act in miRNA maturation, splicing, and ribosome biogenesis pathways, respectively. Together, these data demonstrate that NP reproducibly and specifically engages a unique subset of host RNAs during infection, especially non-coding RNAs transcribed by Pol III.

### Infection regulates the RNA-binding repertoire of NP

Approximately 25% of all NP:RNA complexes contained host RNAs. To determine if these complexes are formed intrinsically or if this is an infection-specific phenomenon, we used spike-in normalized CLIP to quantitatively profile NP-bound RNAs in multiple conditions (Fig S1D-E). NP was expressed via infection, induction in a stable cell line, or both prior to quantitative CLIP-Seq (Fig 3A). Upon infection, ∼30% of NP CLIP reads were influenza virus-derived in infected cells, whereas ∼20% were influenza virus-derived when NP was pre-expressed before infection (Fig 3B, Table S6). Principle coordinate analysis revealed that NP binds a distinct collection of RNAs compared to the input, and this population differs when NP is expressed alone compared to infection (Fig 3C). Over 40% of the identified peaks overlapped between both infection conditions, but less than 15% overlapped between infected cells and those where NP was only present due to inducible expression (Fig 3D). In addition, 3 times more peaks were identified in the absence of infection, suggesting widespread changes in RNA binding dependent on infection. Gene-level analysis of the RNAs bound by NP reflected this differential binding (Table S7). For example, U6 snRNA, one of the most abundant RNAs bound by NP during infection, does not interact with NP outside of infection (Fig 3E, top). vtRNAs and Y4 RNA were also selectively enriched in infection conditions (Fig S5A). Y1 (RNY1) was bound in all conditions with peak intensity increasing from induced to infection settings (Fig 3E, middle). In contrast, 7SK RNA was bound similarly in induced and infected settings (Fig 3E bottom). For both Y1 and 7SK, binding increased and appeared additive when NP was expressed by both infection and induction. Expression of the non-coding RNAs bound by NP remained largely unchanged across all conditions, eliminating the possibility that binding discrepancies are a function of changes in RNA abundance (Fig 3F). Altogether, >1400 peaks are differentially bound between induced and infected samples indicating that influenza virus infection reshapes the population of RNAs bound by NP (Table S6).

**Figure 3.**
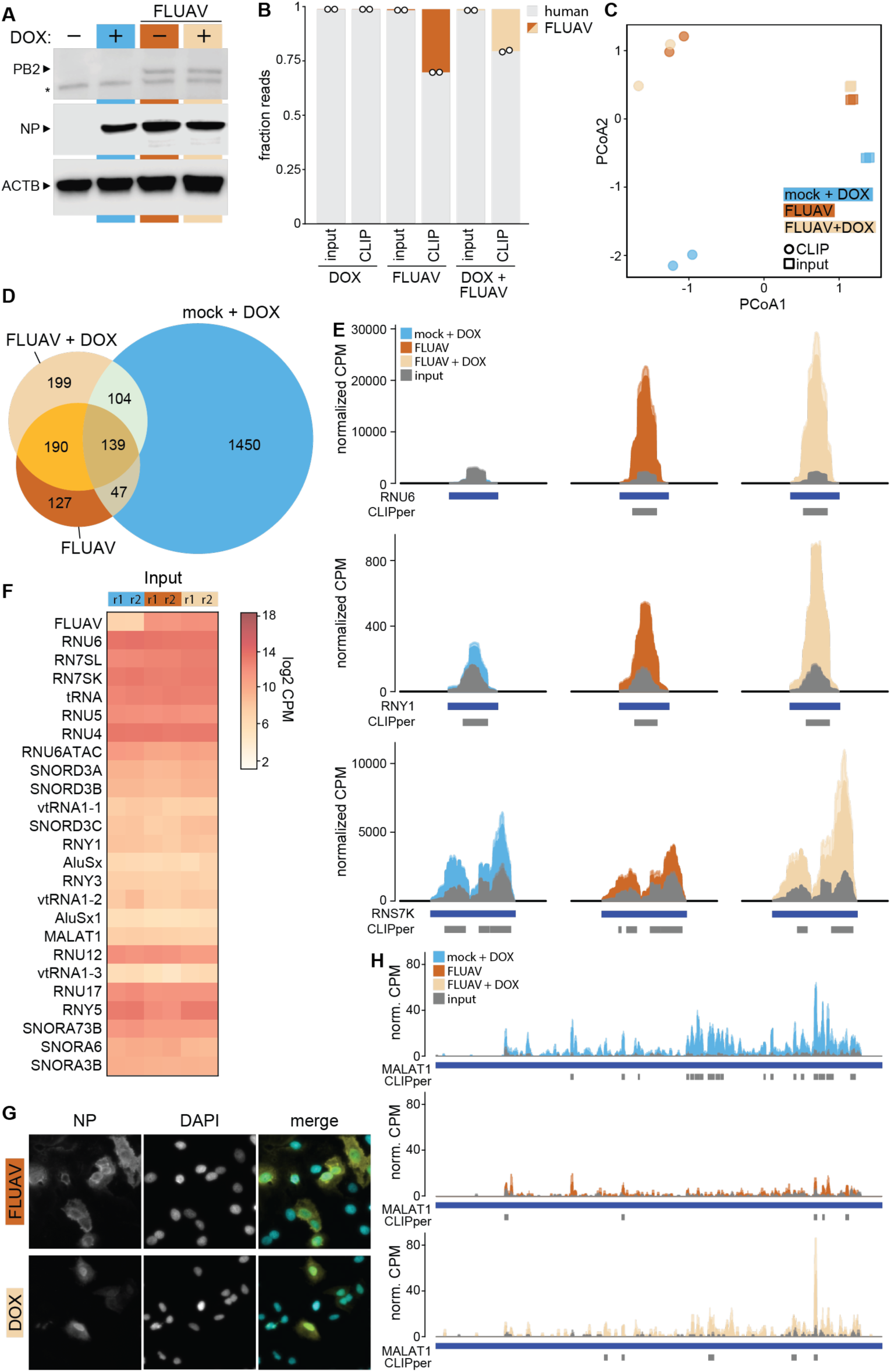
NP RNA-binding is dynamically controlled by infection. (A) Expression of NP in tetNP-A549 cells via doxycycline induction, infection with FLUAV (MOI = 0.01. 24 h), or both. Proteins were detected by western blotting. PB2 served as a control for successful infection control. (B) NP CLIP was performed for the indicated conditions using a shared spike-in control. Frequency of human- and FLUAV-derived reads was determined for NP CLIP and size-matched input control libraries. (C) Principal coordinate analysis of spike-in-normalized NP CLIP and size-matched input control libraries. (D) Overlap of NP-binding peaks for each condition. (E) Normalized read coverage for NP-bound and size-matched input control RNAs. Data from independent biological replicates are shown as mean and s.d. using dark and light shades of the same color, respectively. CLIPper-identified peaks and gene tracks are shown. CPM, counts per million. Associated with Supplemental Tables 6-7. (F). Log2 CPM of select non-coding RNAs in the size-matched input control. (G) Immunofluorescence microscopy of tetNP-A549s stained for NP 24 h after infection (MOI = 0.01) or doxycycline induction. (H) Read coverage as in (E) over the MALAT1 gene body.

The localization of NP is dynamically regulated during infection, beginning in the nucleus early in infection and shifting globally to the cytosol late in infection^47^. Similar patterns are observed when NP is expressed via transfection, although the timing is delayed relative to infection^48^. Immunofluorescence staining confirmed NP follows similar patterns in our cells, with NP globally distributed at the later stages of infection in our experiments while it remains enriched in the nucleus following induced expression (Fig 3G). To test whether the subcellular localization dictates the RNAs that are available and bound by NP, we performed CLIP-Seq using cytosolic and nuclear fractions from infected cells (Fig S5B). Differential binding revealed the NP-bound RNAs enriched during infection over mock correlate with those bound by cytoplasmic NP (Fig S5C). Conversely, MALAT1, an exclusively nuclear-localized lncRNA^49^, is preferentially bound by NP when it is expressed via induction (Fig 3H). These data suggest that the population of host RNAs bound by NP is regulated in part by the subcellular localization of NP, and that NP:RNA interactions may be dynamically regulated throughout the cycle of infection as NP localization changes or other infection-induced factors impact available RNAs.

### NP binds host ncRNAs that signal through RIG-I

RIG-I and NP bound many of the same host transcripts and interacted with each other in an RNA-dependent manner (Fig 2B, 4A, S5E). RNA binding sites for RIG-I and NP showed significant overlap, as demonstrated on 7SL2 RNA (RN7SL2), a component of the signal recognition complex (Fig 4B). Many of the shared RNAs are produced by Pol III, whose nascent transcripts frequently possess hallmarks of immunogenic RNAs sensed by RIG-I including dsRNA termini with a 5’ PPP, although the 5’ PPP is normally processed to a monophosphate to prevent sensing of host transcripts (Fig 4C). Dysregulated sensing of Pol III transcripts is associated with disease^50^. We therefore determined if NP binds immunogenic host RNAs. NP:RNA complexes were immunopurified from infected cells. The immunogenicity of RNAs extracted from these complexes was measured in interferon-stimulated response element (ISRE) reporter assays. These were performed by introducing RNA in human A549 lung cells stably encoding an ISRE reporter that we showed responds to IFN β treatment (Fig 4D). NP-bound RNAs isolated from infected cells dramatically upregulated ISRE activity compared to background activity of RNA extracted from control immunoprecipitations (Fig 4D). To determine if this was a generic activity of NP or required infection, we repeated this experiment using cells where NP was expressed by transfection. Surprisingly, RNA bound by NP purified from transfected cells did not activate the ISRE (Fig 4D). There was significant overlap in the RNAs bound in infected and transfected cells (Fig 3D), but we cannot exclude compositional differences in viral or host RNAs that contribute to differences in immunogenicity.

**Figure 4.**
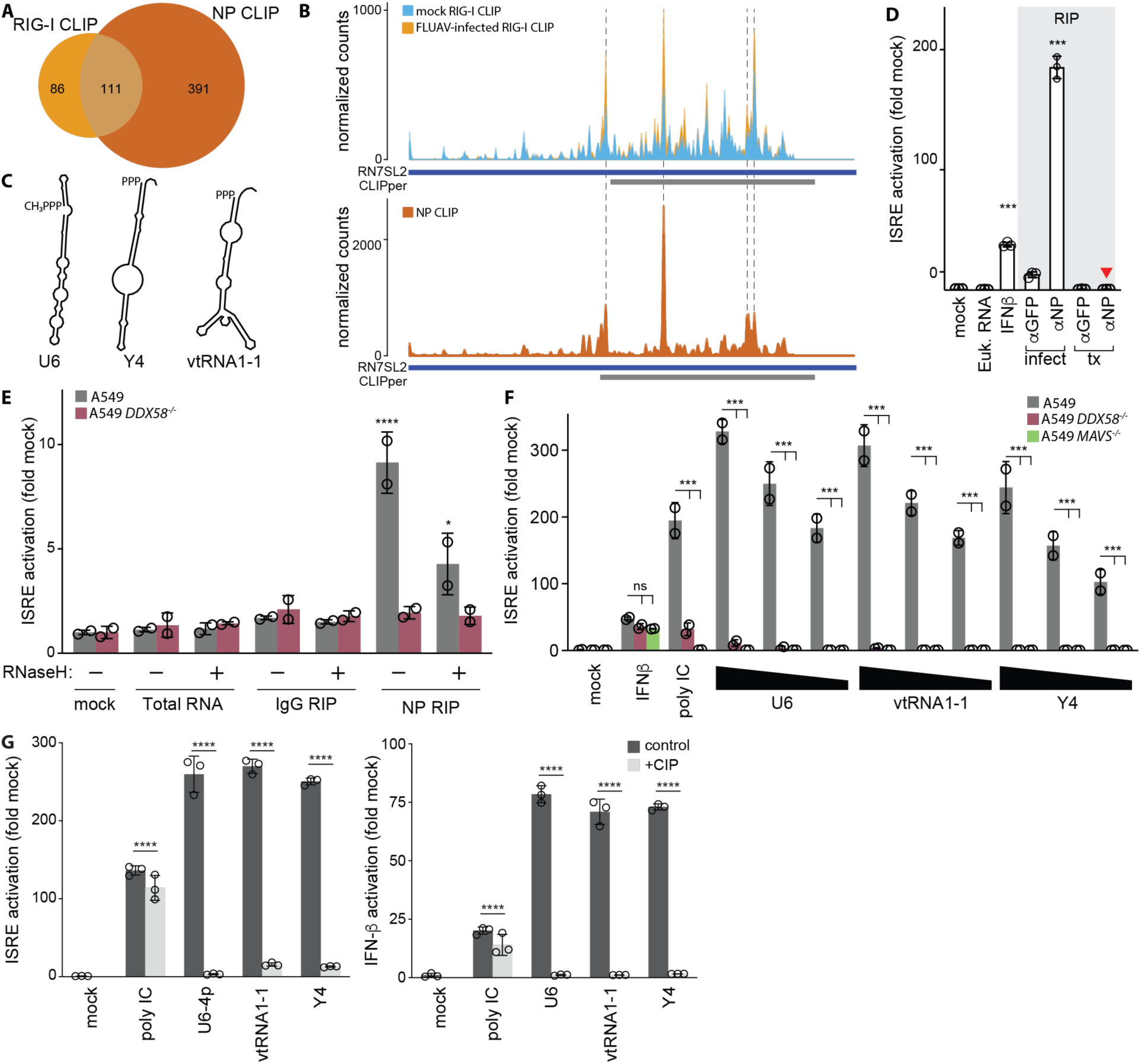
NP-bound non-coding RNAs signal through RIG-I. (A) Overlap of the union of RIG-I CLIP peaks identified in all conditions in in Fig 1 with NP CLIP peaks identified in infected cells in Fig 2E. (B) Read coverage on 7SL RNA of the 5’-mapped base in RIG-I (top) and NP (bottom) CLIP libraries. Coverage was normalized in each library to enable direct comparisons across samples. Data from independent biological replicates are shown as mean and s.d. using dark and light shades of the same color, respectively. CLIPper-identified peaks and gene tracks are shown. (C) Minimum free energy RNA secondary structures predictions from RNAFold for select non-coding RNAs. (D) RNA was isolated from RNA immunoprecipitations (RIPs) performed with the indicated antibodies in cell lysates where NP was expressed by infection (infect) or transfection (tx). 293 ISRE reporter cells were transfected with the recovered RNAs and ISRE activation was measured relative to mock-treated cells. Total eukaryotic RNA or IFN-β were included as controls. Data normalized to an internal *Renilla* control and plotted relative to mock-treated cells. Red arrowhead = lack of activation by RNAs bound by NP in transfected cells. (E) Eluates of NP RNA-immunoprecipitations from FLUAV infected A549 cells (MOI = 0.02, 24h) were treated with an RNaseH-based depletion targeting influenza RNA or mock treated. Remaining RNAs were transfected into WT or RIG knockout (*DDX58^-/-^)* A549 ISRE-reporter cells. ISRE-promoter induction was normalized to an internal *Renilla* control and shown relative to mock-treated conditions for each cell line. (F) WT, RIG knockout (*DDX58^-/-^)*, or MAVS knockout (*MAVS^-/-^*) A549 ISRE-reporter cells were transfected with decreasing amounts of *in vitro* transcribed non-coding RNAs, mixed molecular weight poly IC, mock transfected, or treated with IFN-β. ISRE-promoter induction shown relative to mock-treated conditions for each cell line. (G) Poly IC or *in vitro* transcribed non-coding RNAs were treated with calf-intestinal phosphatase (CIP) or mock treated and transfected into ISRE-reporter cells. Cells were assayed for ISRE-induction and normalized to mock-transfected cells. * P < 0.05, *** P < 0.001, **** P < 0.0001 with one-way ANOVA and Dunnet’s *post-hoc* correction for (D) and a two-way ANOVA with Tukey’s multiple comparison test in (E-G).

NP binds the viral genome and other replication products that are known to activate innate immune responses, explaining some of the observed activation^15–19^. To determine if host RNAs also contributed to innate immune activation, we developed an RNaseH-based depletion strategy that selectively degrades viral RNAs and eliminates their immunogenicity (Fig S6A-B). NP was again purified from infected cells and the associated RNAs were mock- or RNaseH-treated prior to performing ISRE activity assays. NP-bound RNAs still activated the ISRE even after removing viral RNAs, indicating that some of the immunogenic RNAs bound by NP are derived from the host cell. ISRE activity assays were repeated in A549 *DDX58^-/-^* cells to determine if RIG-I senses NP-bound RNAs. Innate immune activation was ablated in the absence of RIG-I for all conditions (Fig 4E). We tested the immunogenicity of individual RNAs by performing ISRE activity assays with pure RNAs created by *in vitro* transcription. U6, vtRNA1-1 and Y4 RNAs each activated the ISRE in a dose-dependent manner (Fig 4F). This activation was lost in cells lacking RIG-I. We additionally tested the role of MAVS, the adaptor that transmits activating signals from RIG-I to downstream effectors. ISRE activation was also lost in *MAVS^-/-^* cells, further implicating RIG-I and its canonical activation pathway in sensing self RNAs. Similar results were obtained when performing activity assays based on the IFN β promoter (Fig S6C). Y RNA and vault RNAs have structured termini with potentially exposed 5’ PPP (Fig 4C). The importance of the 5’ PPP was measured to further establish the role of RIG-I. ISRE activity assays were performed with RNAs treated with calf intestinal phosphatase (CIP) to remove the 5’ PPP. Whereas control samples were highly immunogenic, removal of the 5’ PPP eliminated activation of both the ISRE and IFN β promoter (Fig 4G). Thus, NP binds immunogenic host RNAs that signal via RIG-I to activate innate immunity and a type I interferon response.

### NP counters the antiviral activity of immunogenic host RNA

NP binds a complex mixture of host RNAs (Fig 2E, S4C). To rigorously test the immunogenicity of specific RNAs isolated from biologically relevant settings, RNAs were purified from infected cells via antisense oligonucleotide (ASO) capture^26^ (Fig 5A). This not only purifies specific RNAs, but it retains the molecular features inherent to these RNA in the infected cell. ASO capture was highly specific for the target RNA, but not other RNAs (Fig 5B). The amount of RNA captured was indistinguishable between infected and mock-treated cells, with the exception of PB2 RNA that is only present in infected cells. Similar amounts of highly purified RNAs were then introduced into our ISRE reporter cells. Host U6, Y4 and vtRNA1-1 purified from infected cells potently stimulated the ISRE reporter and one driven by the IFN β promoter (Fig 5C). As expected, genomic PB2 RNA also activated the ISRE and IFN β promoters, whereas non-immunogenic snoRNAs or the non-targeting control capture did not. Importantly, the same RNAs captured from mock-infected cells did not activate innate immune sensing, further supporting the hypothesis that infection-specific events change the immunogenicity of select host RNAs.

**Figure 5:**
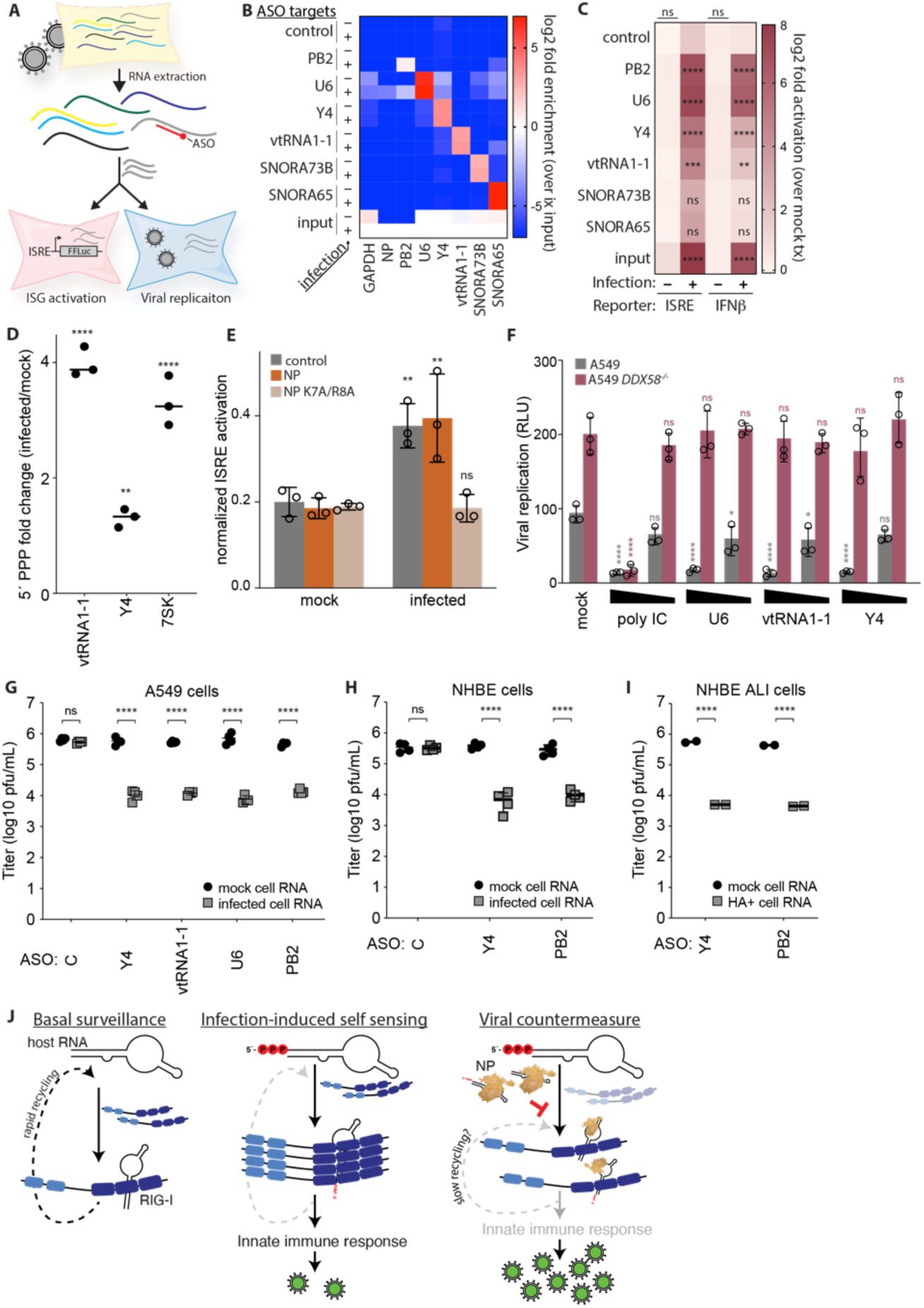
NP counters the antiviral activity of immunogenic host non-coding RNA. (A) Experimental scheme using antisense oligos (ASO) to purify target RNAs prior to re-introduction into cells to measure innate immune activation or viral replication. (B) The indicated RNAs were ASO purified from total RNA extracted from infected or mock-treated A549 cells. The heatmap of qRT-PCR results shows highly specific enrichment of the desired RNAs. Enrichment is shown relative to RNA levels in the input from infected cells. (C) A549 ISRE- or IFN-β-reporter cell lines were transfected with ASO-purified RNAs from mock-treated or infected A549 cells. Induction is relative to mock-transfected cells for each condition. Statistical significance was assessed by comparison to control ASOs in each condition. There were no significant differences when RNAs were purified from uninfected cells. (D) 5’ PPP RNAs were marked by vaccinia capping enzyme, purified, and quantified by RT-qPCR. The fold change in 5’ PPP RNA abudnace was calculated for RNAs from in influenza virus-infected A549 cells relative to those in mock-treated cells. (E) NP or the NLS-mutant NP K7A/R8A was expressed in 293T ISRE-firefly reporter cells for 24 h prior to transfection with Y4 RNA or mock-transfection. ISRE induction was measured relative to an internal *Renilla* luciferase control. (F) WT or RIG-I knockout (*DDX58^-/-^*) A549 cells were transfected with the decreasing concentrations of *in vitro* transcribed non-coding RNA, poly IC, or mock transfected. Cells were subsequently infected with an A/WSN/33-based reporter virus. Supernatants were recovered 24 hpi and titered on MDCK cells. (G) RNA was extracted from mock-treated or infected (A/WSN/33 MOI = 0.1) A549 cells. ASO-purified RNAs were reintroduced into A549 ISRE-reporter cells prior to infection with A/WSN/33 (MOI = 0.5). Viral titers were measured 24 hpi by plaque assay. (H-I) RNA was extracted from normal human bronchial epithelial (NHBE) cells infected with a primary influenza A isolates (A/California/04/2009 MOI = 0.5). NHBE cells in (I) were differentiated at the air-liquid interface (ALI) prior to infection. ASO-purified RNAs were reintroduced into A549 ISRE-reporter cells prior to infection with A/WSN/33 (MOI = 0.5). Viral titers were measured 24 hpi by plaque assay. (J) A model for viral antagonism of self sensing. Under basal conditions, RIG-I rapidly samples and recycles off of non-immunogenic host RNAs maintaining a pool of RIG-I that surveils the entire RNA landscape. During infection, immunogenic features on host RNAs like the presence of a 5’ PPP promote RIG-I oligomerization and activation of innate immune pathways that suppress viral infection. Influenza virus NP counters this by binding host RNAs by interfering with RIG-I activation, possibly by disrupting recycling, to promote viral replication. * P<0.05, ** P < 0.01, *** P <0.001, and **** P < 0.0001 as determined by one-way ANOVA (C,D) with *post hoc* Tukey’s multiple comparisons test or two-way ANOVA (E-I) with *post hoc* Šídák’s multiple comparisons test. Unless noted, comparisons were made to mock-treated cells in each condition. ns = not significant. Note that data in (I) representative data from two independent biological replicates.

Given the importance of the 5’ PPP in activating RIG-I (Fig 4G), we measured changes in 5’ end status during infection. We exploited the fact that vaccinia capping enzyme exclusively modifies RNAs with 5’ PP and PPP moieties^51^. Vaccinia capping enzyme was used to mark these RNAs with a biotinylated cap, which was subsequently used for purification and quantification of the modified RNAs. The change in 5’PP/PPP species for specific RNAs was then quantified by RT-qPCR. Viral infection caused a significant increase in the percentage of vtRNA1-1, Y4 RNA, and 7SK RNA that retained a 5’ PPP relative to mock cells (Fig 5D). The smaller change for Y4 compared to vtRNA1-1 and 7SK resulted from a higher baseline level of 5’ PPP in uninfected cells. This change in the frequency of 5’ PPP likely contributes to the differential immunogenicity of host RNAs in infected cells.

NP binds to immunogenic RNAs that are sensed by RIG-I, raising the possibility that NP may interfere with proper monitoring of the RNA pool. We sought to directly test whether NP interferes with RIG-I sensing. This is not possible in infected cells, as NP and its RNA binding capacity are essential for viral replication and any manipulation will have confounding effects. Instead, we performed ISRE activity assays in the presence of NP. We expressed WT NP that localizes to the nucleus mirroring early stages of infection. We also expressed an NP variant K7A/R8A that localizes to the cytoplasm, the same subcellular compartment as RIG-I, mimicking later stages of infection^47^. WT and NP K7A/R8A displayed the expected subcellular localization (Fig S6D). Pre-expressing WT NP had no effect on the sensing of immunogenic Y4 RNA as ISRE activation was similar to control cells without NP (Fig 5E). However, NP K7A/R8A completely abolished sensing with ISRE activation remaining at the level of mock-treated cells. Thus, cytoplasmic NP prevented the proper recognition and sensing of immunogenic self RNAs.

Sensing self RNAs activates innate immunity. We tested whether this helps establish an antiviral state. Viral replication was measured in A549 lung cells previously transfected with *in vitro*-transcribed self RNAs or poly IC as a control. Viral replication decreased sharply in a dose-dependent manner in cells transfected with self RNA compared to intreated cells (Fig 5F). The magnitude of decrease was dependent on the amount of self RNA the cells received. Infections were repeated in cells lacking RIG-I. Not only did viral replication increase in the absence of RIG-I, self RNAs had no impact on viral replication (Fig 5F). This contrasts with high levels of poly IC that still suppressed replication, indicating that other innate sensing pathways remained active in these cells, suggesting that self sensing of Pol III transcripts is exclusive to the RIG-I pathway.

To reinforce the biological relevance of our results, we tested the antiviral activity of endogenous self RNAs purified from infected cells (Fig 5A). Total RNA was extracted from A549 cells infected with the lab strain A/WSN/33 or mock treated. Target RNAs were then purified with ASOs and transfected into A549 ISRE reporter cells prior to their infection with influenza virus. Introduction of Y4, vtRNA1-1 or U6 RNAs purified from infected A549 cells reduced viral titers by ∼100 fold, but these RNAs had no effect when they were purified from mock-treated cells (Fig 5G). Similar results were obtained when the viral PB2 RNA was purified. In contrast, the non-targeting control purification did not alter replication regardless of the source of the RNA. We then repeated these experiments using RNA purified from primary normal human bronchial epithelial cells (NHBE) that were infected with the pandemic isolate or mock treated. Again, introduction of the self RNA Y4 dramatically reduced viral when purified from infected-cell RNA, but not mock-treated-cell RNA (Fig 5H). The viral RNA PB2 also inhibited replication when purified from infected cell RNA, whereas the non-targeting control did not. Finally, we differentiated NHBE cells at the air-liquid interface (ALI) to create a pseudostratified cell layer that mimics the epithelium of the respiratory tract and infected them with A/California/04/2009. As only ∼20% of the cells in ALI cultures become infected^52^, we first isolated infected cells by flow cytometry prior to extracting the total RNA used for ASO purification. Introducing Y4 RNA purified from infected ALI cells caused a marked reduction in viral replication compared to RNA purified from mock-treated cultures (Fig 5I). Purified viral PB2 RNA showed a similar pattern. The reduction in viral titer caused by RNAs isolated from primary cells correlated with the ability of these self RNAs to activate innate immunity (Fig S7A-B). Together, these data show that self RNAs become immunogenic during infections in biologically relevant settings and contribute to innate immune activation that suppresses viral replication, a process thwarted by NP.

## Discussion

RIG-I patrols the cytoplasm monitoring the RNA landscape for the presence of immunogenic RNAs. This process requires rapid discrimination between viral and host RNAs. Our newly developed quantitative approaches revealed that the majority of RNAs bound by RIG-I are host ncRNAs, even during infection. These host RNAs are benign under basal conditions, but become immunogenic and signal through RIG-I during infection to amplify innate immune responses that suppress influenza virus replication. While self sensing by RIG-I is frequently associated with interferonopathies and chronic inflammatory diseases^4^, it serves a protective function during influenza virus infection. This antiviral process is countered by influenza virus NP, which binds many of these same RNAs and disrupts RIG-I activation. Together, these data show that self sensing is a key component of the antiviral response to influenza virus and identify a viral countermeasure that directly disrupts sensing of host ncRNAs.

RIG-I surveils a vast excess of RNAs as it seeks the rare specimen that is immunogenic. The baseline binding we identified in uninfected cells represents RIG-I actively sampling and recycling. Our quantitative approaches revealed that RIG-I does not preferentially bind viral RNAs, fitting well with kinetic models where RIG-I discriminates immunogenicity not at the association stage, but through altered dissociation rates^12,22,53^. This allows for rapid recognition of immunogenic RNAs when they are introduced into the system, whether from the virus or the host. For example, host Y4, U6, and vtRNA1-1 all activated RIG-I sensing, but only when purified from infected cells (Fig 5). This suggests that the immunogenicity of small ncRNAs is differentially regulated during infection. Almost all of the host RNAs bound by RIG-I are non-coding RNAs transcribed by RNA Pol III, whose products initially contain a 5’ PPP^54^. Previous work identified the dual-specificity phosphatase 11 (DUSP11) as a regulator of self sensing where it prunes phosphates from 7SL, RNY4, and vtRNA1-1 to prevent activation of RIG-I^26,29,55^. Our data show that 5’ end status also changes during FLUAV infection to enable self sensing, implicating retention of 5’ PPP on host RNAs as a common strategy to amplify innate immune activity across divergent viral infections (Fig 5J). This reinforces an emerging theme where cells deploy host RNA or DNA during infection to strategically activate self sensing^25–28,55^.

RNA viruses frequently encode proteins that disable RLRs^56,57^. This inhibition can be mediated by direct interaction with RLRs, disruption of co-factors that are required for RLR activation, interception of signaling cascades downstream of RLRs, or by directly binding viral RNAs to prevent sensing^57^. The FLUAV protein NS1 is a primary RIG-I antagonist that suppresses antiviral responses by binding viral dsRNA and targeting cellular co-factors needed for RIG-I activation^58–60^. Whether NS1 alters the repertoire of RNAs bound by RIG-I or NP remains to be determined. NP is also thought to prevent sensing of viral genomic material by encapsidating viral RNAs in RNPs^16^. Our data demonstrate a previously unappreciated role for NP and identify a new strategy to specifically disrupt self sensing by binding host ncRNAs. There are several possible mechanisms by which NP could disrupt RIG-I activation. Given its high expression and non-specific RNA-binding properties^41,46^, NP may simply bind and sequester host ncRNAs from RIG-I. NP also melts dsRNA^40^, potentially eliminating key features in the RNAs needed for RIG-I recognition. The outcome of both of these would be lower levels of host RNAs bound by RIG-I. However, this is not seen as the abundance of host ncRNAs on RIG-I is indistinguishable between mock and infected conditions (Fig S2C). Instead, we speculate that NP may slow RIG-I recycling (Fig 5F). NP and RIG-I interact in an RNA-dependent manner binding the same RNAs in the same regions (Fig 2B, 4B). If NP slows RIG-I recycling, whether on benign or immunogenic RNAs, this will rapidly deplete the pool of free RIG-I and prevent formation of activated oligomers, even if RIG-I ultimately bound immunogenic RNAs. This possibility would also mean that RIG-I antagonism is not restricted to interactions on the small number of host RNAs that become immunogenic, but any RNA occupied by both NP and RIG-I. There is precedent for RIG-I interacting with viral proteins during translocation on dsRNA^61^. Whether this results in longer retention times on non-immunogenic RNAs remains to be determined.

All negative-strand RNA viruses, and some positive-strand RNA viruses like coronavirus, express NP or an N protein that oligomerizes on viral RNA in a sequence agnostic fashion^35,62,63^. NP/N is expressed at high levels to facilitate genome replication. These levels far exceed the quantity present in RNPs and thus hint at the existence of additional roles for nucleoproteins^64–66^. The shared features of NP/N proteins suggest that the properties we defined here for FLUAV NP may be broadly relevant to a large swath of RNA viruses. Sequence-independent associations between NP/N and host RNAs may provide a universal and remarkably simple strategy to disrupt self sensing by RIG-that tempers innate immune responses to promote replication.

## Data Availability

Raw data are available in Table S9 and uncropped blots are available in Fig S8. Sequencing data have been deposited at BioProject PRJNA1213036, PRJNA1213052, PRJNA1213053. Sequencing statistics and accession numbers are summarized in Table S8.

## Acknowledgements

This work was supported by the National Institutes of Health R01AI164690, R21AI125897, UW2020 Initiative by the Wisconsin Alumni Research Foundation, and the Tracy, Dawn and Jamie Ruhrup Memorial Fund awards to AM and the Canadian Institutes of Health Research (CIHR) Project Grant PJT 148727 to DAK. MPL was supported by a National Science Foundation grant GRFP DGE-1747503 and TN by T32AI007414. AM is an H.I. Romnes Faculty Fellow funded by the Wisconsin Alumni Research Foundation, a Vilas Faculty Mid-Career Investigator, and completed portions of this work as a Burroughs Wellcome Fund Investigator in the Pathogenesis of Infectious Disease. We thank members of the Mehle laboratory for valuable input and M. Harrison and N. Grandvaux for the gifts of reagents.

## Author Contributions

Conceptualization – MPL, AM

Data curation – MPL, AM

Formal analysis – MPL, AM

Funding acquisition – MPL, TN, AM

Investigation – MPL, TN, KAD, DU, ZZ

Methodology – MPL, TN, KAD, DU, ZZ, MK, DAK

Project administration – AM

Resources – MPL, TN, KAD, AM

Software – MPL, TN

Supervision – DAK, CM, AVK, AM

Validation – MPL, AM

Visualization – MPL, DU, AVK, AM

Writing – original draft – MPL, AM

Writing – review & editing – MPL, TN, KAD, DU, AVK, AM

## Declaration of Interests

The authors declare no known conflicts of interest.

## SUPPLEMENTAL INFORMATION

**Document S1.** Figures S1–S8

**Supplementary Table 1:** RIG-I-bound RNAs. Associated with Fig 1.

**Supplementary Table 2:** Differential binding of RIG-I to RNAs in mock and infected cells. Associated with Fig 1.

**Supplementary Table 3:** Differential binding of RIG-I at CLIP peaks in mock and infected cells. Associated with Fig 1.

**Supplementary Table 4:** Differential binding of NP to RNAs in infected cells analyzed at the gene level. Associated with Fig 2.

**Supplementary Table 5:** NP-bound RNAs in infected cells. Associated with Fig 2.

**Supplementary Table 6:** NP-bound RNAs during induction or infection. Associated with Fig 3.

**Supplementary Table 7:** Differential binding of NP to RNAs in infected or doxycycline-induced cells. Associated with Fig 3 and S4.

**Supplementary Table 8:** Statistics and accession numbers for sequencing data

**Supplementary Table 9:** Raw data from figures

## Methods

### Cells

Human lung alveolar epithelial (A549) cells (CCL-185), Madin-Darby canine kidney (MDCK) cells (CCL-34), baby hamster kidney-21 (BHK-21) cells (CCL-10), and human embryonic 293T (HEK293T) cells (CRL-3216) were purchased from the American Type Culture Collection (ATCC) and maintained in Dulbecco’s modified Eagle’s medium (DMEM) and 10% fetal bovine serum (FBS) at 37°C and 5% CO_2_. *Drosophila* Schneider 2 (S2) cells were maintained in Schneider’s *Drosophila* medium supplemented with 10% FBS and 1% antibiotic/antimycotic and grown at 25°C without CO_2_. A549*DDX58^-/-^* cells were prepared by CRISPR/Cas9-mediated mutagenesis. A549 *MAVS^-/-^* cells were a gift from N. Grandvaux (Université de Montréal)^67^. Both knockout cell lines were used with their parental wild-type A549 cells. HEK293 cells stably expressing N-terminally STrEP-tagged RIG-I or mCherry were previously described^68^. Primary normal human bronchial epithelial cells were from a Caucasian white male age 55. Cultures were expanded in PneumaCult™-Ex Plus Medium (Stem Cell Technologies #5040) and differentiated for >21 days on collagen-coated transwells at the air-liquid interface prior to infection using the PneumaCult-ALI medium system (Stem Cell Technologies Cat# 05001). Cells were regularly verified to be mycoplasma negative using MycoAlert (Lonza).

### Virus

Influenza virus A/WSN/33, A/WSN/33-based PA-2A-Nanoluc reporter virus, and A/California/04/2009 were grown and titered as described befored^69–71^. In brief, viral stocks were grown in MDCK cells and titered on MDCK cells with an Avicel overlay. Vesicular stomatitis Indiana virus encoding GFP was grown and titered by plaque assay with agarose overlay using BHK-21 cells.

### Antibodies

The following antibodies were used as indicated: H16-L10-4R5 (HB-65) mouse monoclonal anti-NP (BioXCell BE0159); B-2 mouse monoclonal anti-GFP (Santa Cruz Biotechnology SC-9996); Normal rabbit IgG (Cell Signaling Technologies 2729); M2 mouse monoclonal anti-FLAG (Sigma F1804); Goat anti-RNP (BEI Resources NR-3133); DM1A mouse monoclonal anti-tubulin (SigmaAldrich T6199); Mouse anti-beta-actin (Proteintech 66009-1-Ig); Rabbit anti-lamin A/C (Cell Signaling Technologies 2032); and, DYKDDDDK Fab-Trap (ChromoTek ffa)

### Plasmids and *in vitro* transcribed RNAs

WT and mutant NP (A/WSN/33) expression plasmids were generated by PCR. Doxycycline-inducible NP plasmids were generated by cloning NP into pSBtet-BP (Addgene 60496) to create pSBtet-BP-NP. The constitutively expressed RIG-I plasmid was created by cloning RIG-I with a 2x FLAG tag into pSBtet-BP to replace BFP creating pSB-RIGIP. The copper-inducible *Drosophila* plasmids pMT-puro-NP and pMT-puro-RIG-I were generated by cloning NP (A/WSN/33) and RIG-I-2xFLAG into pMT-puro (Addgene 17923). The ISRE- and IFN-β -response element plasmids pSB-BB-ISRE-NLuc2AG and pSB-BB-IFNβ-NLuc2AG encode a NanoLuc-2A-GFP fusion protein driven by an ISRE promoter or IFN-β promoter, respectively, on the pSBbi-BB plasmid backbone (Addgene 60521). The ISRE-response element plasmid pLX304-ISRE-FFLuc was generated by cloning the ISRE-FFLuc cassette into pLX304 (Addgene 25890).

Non-coding RNAs were reverse-transcribed from total RNA with gene-specific primers and cloned via Gibson assembly into a plasmid for maintenance. These plasmids were used in PCRs to create templates for T7 RNA polymerase transcription. Non-coding RNA genes were amplified with gene-specific primers that appended a T7 RNA polymerase promoter on the 5’ end. The product was purified and 2 µg DNA template was *in vitro* transcribed in a 100 μl reaction with 40 mM Tris pH 7.9, 6 mM MgCl_2_, 10 mM DTT, 2 mM spermidine, 2mM rNTP, 1 μl RNasin(+) (Promega N2611), and 3 μl Hi-T7 RNA polymerase mix (NEB M0658S) for 2h at 37°C. RQ1 DNaseI (Promega M6101) was added and the reaction was incubated for 30 minutes at 37°C. RNA was purified via phenol-chloroform or TRIzol extraction and the resultant product checked for purity and size via TBE denaturing urea polyacrylamide gel electrophoresis and stained with SYBR gold (Invitrogen). The following RNAs were produced:

RNY4: ggcugguccgaugguaguggguuaucagaacuuauuaacauuagugucacuaaaguugguauacaaccccccacugcuaaauuugacuggcuuuuu

RNU6: gugcucgcuucggcagcacauauacuaaaauuggaacgauacagagaagauuagcauggccccugcgcaaggaugacacgcaaauucgugaagcguuccauauuuuu

vtRNA1-1: ggcuggcuuuagcucagcgguuacuucgacaguucuuuaauugaaacaagcaaccugucuggguuguucgagacccgcgggcgcucuccaguccuuuuu

To remove the 5’-triphosphate from RNAs, 2 µg of *in vitro* transcribed RNAs were mock-incubated or treated with calf-intestinal phosphatase. RNA was purified via RNA clean-and-concentrator kits (Zymo Research R1014).

### Generation of transduced cell lines

Vesicular stomatitis Indiana virus (VSIV) glycoprotein G-pseudotyped lentivirus was generated by transfecting 293T cells with the plasmids psPAX2 and pMD2.G, along with pLX304-ISRE-FFLuc. The resultant lentivirus was inoculated on A549 cells to create firefly luciferase ISRE reporter cells. Cells were allowed to recover and selected with 6 µg/mL blasticidin. NanoLuc ISRE- and IFN-β reporter cells were generated with the Sleeping Beauty transposon system by transfecting A549 cells with equal amounts of pSB-ISRE-NLuc2AGFP or pSB-IFNβ-NLuc2AGFP and pCMV(CAT)T7-SB100, a plasmid expressing the Sleeping Beauty transposase (Addgene 34879)^72^. Cells were allowed to recover and selected in 6 µg/mL blasticidin. Doxycycline-inducible NP A549s and stable RIG-I expressing A549s were generated similarly, except pSBtet-BP-NP and pSB-RIG-I were transfected with the transposase and selected in 0.5 µg/mL puromycin. Copper inducible NP and RIG-I S2 cells were generated by transfection of pMT-puro-NP and pMT-puro-RIG-I with Transit-X2 (Mirus) and selection with 2 µg/mL puromycin for 2-4 weeks.

### ISRE and IFN-β reporter assays

Cells were transfected with *in vitro* transcribed RNA, ASO-purified RNA (see below), poly IC, or mock transfected using Transit-X2 (Mirus Bio). Cells were incubated for 24h, lysed in *Renilla* Luciferase Assay buffer (Promega), and assayed for NanoLuc, firefly luciferase, and/or *Renilla* luciferase depending on the reporter system used. In some experiments, cells were imaged for GFP expression.

For NP/RNA co-transfection assays, 293Ts were plated in 96-well plates and transfected using Transit-X2 (Mirus Bio) with 90 ng control plasmid, pEF1-NP-3xFLAG, pEF1-NP-K7AR8A-3xFLAG, and 10 ng pRL-SV40 and incubated for 24 hours. Immunogenic RNA was introduced at 25 ng/well via transfection with Transit-X2 (Mirus Bio). Cells were incubated for 24 h and assayed for firefly and *Renilla* luciferase expression. Data were normalized to the *Renilla* control.

### RNA immunoprecipitations (RIPs)

A549 or A549-RIG-I cells were infected with A/WSN/33 (MOI = 0.2, 24h). NP RIPs were lysed in co-immunoprecipitation buffer (50 mM Tris pH 7,5, 150 mM NaCl, 0.5% NP-40) while RIG-I RIPs were lysed in RIG-I RIP buffer (20 mM Tris–HCl (pH 7.5), 150 mM NaCl, 0.25 % NP-40, 1.5 mM MgCl2, 1 mM NaF), both supplemented with 400 U/mL RNasin (Promega) and protease inhibitor cocktail (Pierce). Lysates were clarified by centrifugation at 15,000 x g for 15 minutes at 4°C and pre-cleared with protein A agarose (Santa Cruz) for 1 h at 4°C. Before immunoprecipitation, RNA was extracted from 10% of input lysate with TRIzol. For NP RIPs, the remaining lysates were incubated with the mouse monoclonal antibodies H16-L10 directed against NP (BioXCell BE0159), the non-specific control B-2 directed against GFP (Santa Cruz sc9996), or normal rabbit IgG (Cell Signaling Technologies 2729). Antibody bound protein-RNA complexes were captured with protein A agarose for 1 h. Protein A beads were washed 5 times with cold co-immunoprecipitation buffer. For RIG-I RIPS, lysates were incubated with DYKDDDDK Fab-Trap (Chromotek ffa) or V5-trap agarose as a generic IgG control (Chromotek v5ta). Trap agarose resin was washed 5 times with cold RIG-I RIP buffer. Following washes, RNA was eluted by resuspending the resin in 100 μl of TRIzol. RNA was then purified using the direct-zol kit (Zymo Research R2052) with included DnaseI digestion. Equal volumes of RNA were utilized for qRT-PCR or sized on an Agilent 2100 Bioanalyzer RNA pico chip.

For qRT-PCR, equivalent volumes of RNA were reverse-transcribed with random-hexamer primed MMLV-RT. The resulting cDNA was diluted 1:10 and used for qPCR with the iTaq Universal SYBR Green Supermix (Bio-rad 1725121) on a Step One Plus RT-PCR instrument (Applied Biosystems) using forward and reverse primers described below. Fold enrichment relative to the input or the IgG control was calculated with the ΔCt method. qPCR primers indicated below.

RNU6_F: GCTTCGGCAGCACATATACTAAAAT; RNU6_R: CGCTTCACGAATTTGCGTGTCAT

RNY4_F: GTCCGATGGTAGTGGGTTATC; RNY4_R: AAAGCCAGTCAAATTTAGCAGT

vtRNA1-1_F: CGACAGTTCTTTAATTGAAACAAGC; vtRNA1-1_R: AAGGACTGGAGAGCGCCC

FLUAV_NA_F: ACATCACTTTGCCGGTATCAGGGT; FLUAV_NA_R: ACCATAATGACCGATGGCCCAAGT

PB2_vRNA_F: CTTTTGGTCGCTGTCTGGCT; PB2_vRNA_R: TGAACTGAGCAACCTTGCGA

RNY4-Fv2: GGCTGGTCCGATGGTAGT; RNY4-Rv2: AAAAGCCAGTCAAATTTAGCAGT

vtRNA1-1-Fv2 :AGCTCAGCGGTTACTTCGAC; vtRNA1-1-Rv2: AAAAGGACTGGAGAGCGCC

RNA18SN1-F: GTTCAGCCACCCGAGATTGA; RNA18SN1-R: CCCATCACGAATGGGGTTCA

RNU6-Fv2: CGCTTCGGCAGCACATATACTA; RNU6-Rv2: TGGAACGCTTCACGAATTTGC

### RNaseH vRNA depletion assay

RnaseH depletion of target RNAs was performed essentially as described, with slight modifications^73^. Briefly, 100 ng of *in vitro* transcribed influenza RNAs or equal volumes of RNAs from NP RNA-immunoprecipitations were diluted into 10 μl total in water with a final concentration of 1 μM dNTPs and 1.25 µM probe mixture containing gene-specific and universal primers targeting the conserved termini of the flu genome (listed below). Samples were heated at 65°C for 5 minutes, and snap-cooled on ice to anneal the probe mixtures to the RNA. A 10 µL mixture containing 2x First Strand Buffer, 20 U Rnasin(+), 10mM DTT, and 0.5 µL SuperScript III reverse transcriptase (ThermoFisher Scientific 18080044) was added to the RNA and incubated at room temperature for 5 minutes, 42°C for 30 minutes, and 50°C for 30 minutes. To digest DNA:RNA hybrid products, 2.5 U RnaseH (ThermoFisher Scientific EN0202) was added to the reverse transcription reaction with 2 μl First Strand Buffer in a 10 µL volume and allowed to incubate at 37°C for 30 minutes. This reaction was mixed with 2 µL RQ1 DnaseI (Promega M6101) in buffer and allowed to incubate at 37°C for 15 minutes. The reaction was terminated by addition of buffer RLT (Qiagen) and purified with Dynabeads MyOne-silane beads (ThermoFisher Scientific 37002D) and eluted in a 5 µL volume of water. Successful depletion of RNA was confirmed via TBE denaturing urea polyacrylamide gel electrophoresis and staining with SYBR-gold. Primers used were:

NS_minu_483-464: CTTCGGTGAAGGCCCTTAGT; NS_plus_440-459: TGACCGGCTGGAGACTCTAA

M_minu_514-495: CCTATGAGACCGATGCTGGG; M_plus_462-481: TGGTATGCGCAACCTGTGAA

NA_minu_1006-987: TCCGTTTGCTCCATCAGCAG; NA_minu_518-499: CACTTGCTGACCAAGCAACC

NA_plus_926-945: AGTGGGGTTTTCGGTGACAA; NA_plus_468-487: CTCCGTCCCCGTACAATTCA

NP_minu_1041-1022: TGCCATCCACACCAGTTGAC; NP_minu_528-509: GGGATCCATTCCTGTGCGAA

NP_plus_1091-1110: GGACGAAAGTGGTCCCAAGA; NP_plus_536-555: GCTCACTGATGCAGGGTTCA

HA_minu_1128-1167: TCATTCTGATGATGATAACC; HA_minu_642-623: GCTCATCACTGCTAGACGGG

HA_plus_530-549: GCTGACGAAGAAGGGGGATT; HA_plus_1153-1172: GATCAGGCTATGCAGCGGAT

PA_minu_516-497: ATCGAGAGTGTAGTCGGCCT; PA_minu_1152-1133: TGGTGCCATGTTCTCACCAA

PA_minu_1697-1678: GACACATGGCCTATGGCACT; PA_plus_1758-1777: GATGGAAATGAGGCGTTGCC

PA_plus_1225-1244: AGGTCGCTTGCAAGTTGGAT; PA_plus_660-679: CAAGCTTGCCGACCAAAGTC

PB1_minu_415-396: AGTCATAGGTCTGTCGGCCT; PB1_minu_848-829: CCTCCAACTGGCAATCCTGA

PB1_minu_1662-1643: CATTTGAGCGGTTGCTGGAC; PB1_plus_1562-1581: GGGTGTCTGGGATCAACGAG

PB1_plus_829-848: TCAGGATTGCCAGTTGGAGG; PB1_plus_375-394: ACGAGTGGACAAGCTGACAC

PB2_minu_2097-2078: AACTGCGGACTCAACTCCAG; PB2_minu_1217-1198: GCTTCGGCAATCGACTGTTC

PB2_minu_676-657: ATCTCGTTTTGCGGACCAGT; PB2_plus_1509-1528: AGTGGTGAGCATTGACCGTT

PB2_plus_1053-1072: AGAGGTGCTTACGGGCAATC; PB2_plus_533-552: CTAACGAAGTGGGAGCCAGG

uni13: AGTAGAAACAAGG; uni12deg: AGCRAAAGCAGG

### Generation of biotin-labeled oligos

Enzymatic synthesis of biotin-labeled capture probes was performed essentially as described^74^. Briefly, RNA complementary to the designed probes was generated by *in vitro* transcription reactions with T7 RNA polymerase using templates created by PCR as described above. Primers were:

capture_RNU6_F: TAATACGACTCACTATAGGgcccctgcgcaaggatga

capture_RNU6_R: ACCAGATCGTTCGAGTCGgaacgcttcacgaatttgcg

capture_RNY4_F: TAATACGACTCACTATAGGgtcactaaagttggtatacaa

capture_RNY4_R: ACCAGATCGTTCGAGTCgaaGcaaatttagcagtgg

capture_vtRNA1-1_F: TAATACGACTCACTATAGGGAAACAAGCAACCTGTCTGGGT

capture_vtRNA1-1_R: ACCAGATCGTTCGAGTCGGCCCGCGGGTCTCGAACAA

capture_PB2_F: TAATACGACTCACTATAGGGTCTGGCTGTCAGTAAGTATGCT

capture_PB2_R: ACCAGATCGTTCGAGTCGAACGGAAACGGAACTCTAGC

capture_SNORA73B_F: TAATACGACTCACTATAGGgcttcccagagtcctgtgga

capture_SNORA73B_R: ACCAGATCGTTCGAGTCGgtttgtctccccagtcattgt

capture_SNORA65_F: TAATACGACTCACTATAGGGTTGTTCTTCATGTGGATGACTC

capture_SNORA65_R: ACCAGATCGTTCGAGTCGCCCATGCTTTCGGCACAG

The resultant RNA was purified by phenol-chloroform extraction and 10 µg was used in a reverse transcription reaction with MMLV and a biotin-tagged reverse transcription primer: biotin-ACCAGATCGTTCGAGTCG. The reaction was terminated after 3 h by hydrolysis of the RNA via addition of 1N NaOH for 15 minutes at 72°C, neutralization of the NaOH by addition of 1N HCl, and purification with a column-based cleanup. The resulting cDNA was checked for size and purity via TBE denaturing urea polyacrylamide gel electrophoresis with SYBR-gold staining and resulted in the probe sequences:

Control ASO: biotin-ACCAGATCGTTCGAGTCG

RNU6 ASO: biotin-ACCAGATCGTTCGAGTCGgaacgcttcacgaatttgcgtgtcatccttgcgcagggg

RNY4 ASO: biotin-ACCAGATCGTTCGAGTCGAAgcaaatttagcagtggggggttgtataccaactttagtgac

vtRNA1-1 ASO: biotin-ACCAGATCGTTCGAGTCGgcccgcgggtctcgaacaacccagacaggttgcttgtttc

PB2 ASO: biotin-ACCAGATCGTTCGAGTCGaacggaaacggaactctagcatacttactgacagccagac

SNORA73B ASO: biotin-ACCAGATCGTTCGAGTCGgtttgtctccccagtcattgtccacaggactctgggaagc

SNORA65 ASO: biotin-ACCAGATCGTTCGAGTCGcccatgctttcggcacagagtcatccacatgaagaacaac

For experiments in Fig 5G-I, biotin-labeled oligos were obtained from IDT.

### Anti-sense oligonucleotide RNA capture

RNA samples for anti-sense oligonucleotide captures (ASOs) were generated by mock-infecting or infecting A549 cells with A/WSN/33 (MOI = 0.02, 24 h) or NHBE cells with A/California/04/2009 (MOI = 0.5, 24 h). Total RNA was extracted with Trizol. 10-25 µg total RNA was mixed with 100 ng DNA probe in binding buffer (10 mM HEPES pH 7.5, 500 mM LiCl, 1 mM EDTA) and incubated at 65°C for 5 m, followed by binding for 15 m at room temperature. 10 µL of Dynabead Streptavidin C1 beads (ThermoFisher Scientific 65002) or 7.5 μl of hydrophilic streptavidin magnetic beads (NEB S1421S) were mixed with oligonucleotide-bound RNAs and incubated with shaking at 25°C for 30 minutes. The beads were magnetically separated and washed with low-salt wash buffer (20 mM Tris pH 7.5, 200 mM LiCl, 1 mM EDTA) 4 times, dried, and resuspended in elution buffer (20 mM Tris pH7.5, 1 mM EDTA). Elution was carried out by heating at 95°C for 1.5 minutes and quickly separating the beads from the supernatant. The following modifications were made when RNA was isolated from ALI cultures infected with A/California/04/2009. Only ∼20% of the cells in an ALI culture become infected^52^, therefore infected cells were first isolated by flow cytometry prior to extraction of total RNA. Only 3 μg of total RNA was available to be used for each replicate. This RNA was subject to serial ASO capture for Y4 followed by PB2. Purified RNAs were used in assays for innate immune activation described above in “ISRE and IFN-β reporter assays” or during infections as detailed in “Multicycle viral growth.”

### Multicycle viral growth

To determine the effects of ASO-purified RNAs on viral replication, A549-ISRE NLuc2AGFP reporter cells were transfected with eluted RNA using Transit-X2 (Mirus Bio) (n=4 for RNA purified from A549 and NHBE cells, n=2 for ALI cultures). Cells were incubated for 24 h, imaged for GFP expression, and infected with A/WSN/33 (MOI = 0.5) in in viral growth medium (DMEM, 0.3% bovine serum albumin, 25 mM HEPES, 1x penicillin-streptomycin, and 0.25 µg/mL TPCK-treated trypsin) for 24 h. Supernatants were harvested and viral titers were measured by plaque assay.

*In vitro* transcribed immunostimulatory RNAs were prepared as described above and transfected into WT or *DDX58^-/-^* A549 cells using Trans-X2 (Mirus Bio). Viral infections were performed 24 h later with a A/WSN/33-based reporter virus expressing NanoLuc as self-cleaving polyprotein with PA^70,71^. Virus was diluted in viral growth medium (DMEM, 0.3% bovine serum albumin, 25 mM HEPES, 1x penicillin-streptomycin, and 0.25 µg/mL TPCK-treated trypsin) at a multiplicity of infection of 0.05. Infections were allowed to progress for 24 h at which point the supernatants were harvested and titered by inoculation of MDCK cells. Titers were measured 16 h later by lysing cells in Renilla Luciferase Assay buffer and assaying for NanoLuc expression^75^.

### eCLIP-sequencing

Unless noted, CLIP-Seq experiments were performed on two independent biological replicates. The NP enhanced CLIP (eCLIP) protocol was previously adapted from published work and performed as described^76–78^. For CLIP-Seq without a spike-in control in Figure 2, 10^7^ A549 cells were infected with A/WSN/33 (MOI = 0.02, 24 h) and washed with PBS immediately before UV-crosslinking with 400 mJ/cm^2^, then 200 mJ/cm^2^ 254 nM UV light after which cells were scraped, collected, and flash frozen. Cells were lysed in CLIP lysis buffer (50 mM HEPES pH 7.5, 150 mM KCl, 2 mM EDTA, 0.5% v/v NP-40, 1 mM DTT, 0.5% DOC w/v, and EDTA-free protease inhibitor). For subcellular fractionation on NP CLIP samples, cells were infected with A/WSN/33 (MOI = 0.01, 24 h), collected on ice, processed via hypotonic lysis to obtain cytoplasmic extracts (n=1), and followed by a high salt extraction to obtain nuclear extracts (n=1). Fraction purity was verified by blotting for tubulin (cytoplasmic) or lamin (nuclear). Lysates were digested with 0.1 U/µL RNaseT1 (ThermoFisher Scientific EN0542) for 10 minutes at room temperature. Size-matched inputs were taken and NP was immunopurified by adding Dynabeads protein A (Invitrogen 10002D) pre-bound with monoclonal NP H16-L10 (BioXCell BE0159) to the lysates and incubated for 3 h at 4°C. Beads were washed twice in lysis buffer, twice in high-salt buffer (50 mM Tris pH 7.4, 1 M NaCl, 1 mM EDTA, 0.1% SDS, 0.5% DOC, 1% NP-40) and twice in lysis buffer followed by labeling the RNA with γ^32^P-ATP. Labeled complexes were separated via SDS-PAGE, transferred to a nitrocellulose membrane, visualized by phosphorimaging, NP-bound reagents were cut from the membrane, and complexes were digested off the membrane with proteinase K. RNA was captured by column purification (Zymo Research R1014) and processed for ligation with a pre-adenylated DNA adapter (New England Biolabs S1315) to generate eCLIP libraries as described^77^.

Spike-in controls were generated in S2 cells. 6x10^7^ copper-inducible NP or RIG-I S2 cells were plated and treated with 500 µM CuCl_2_ for 48 h. Cells were collected, cross-linked, and lysed before digestion with 0.05 U/uL RNaseT1 (ThermoFisher Scientific EN0542) for 10 minutes at room temperature. Lysates were aliquoted and flash frozen.

A549 eCLIP libraries were prepared with a spike-in control. For A549-tetNP cells, 10^7^ cells were either treated with 2.5 µg/mL doxycycline, infected with A/WSN/33 (MOI = 0.01, 24 h), or both infected and treated. For A549-RIG-I cells, 10^7^ cells were infected with A/WSN/33 (MOI = 0.01, 24 h), infected with VSV-GFP (MOI = 0.001, 24 h), or mock-infected. In all cases, cells were washed with cold PBS, cross-linked, and flash frozen. Lysates were processed as described above with the following exceptions. Lysates were treated with 0.05 U/uL RNaseI (ThermoFishewr AM2294) for 10 minutes at room temperature, quenched by transfer to ice, and a shared spike-in control was added by mixing each sample with the same amount of *Drosophila* lysate at ∼1:1 A260 OD ratio. Lysates were immunoprecipitated with monoclonal NP H16-L10 (BioXCell BE0159) or monoclonal M2 (Sigma F1804) bound to protein A dynabeads, dependent on the target RNA-binding protein. Beads were washed twice with lysis buffer, twice with high salt buffer, and twice with lysis buffer before separation via SDS-PAGE and transfer to a nitrocellulose membrane. RNA samples were processed as above, except samples were ligated with the pre-adenylated DNA linker: AppAGATCGGAAGAGCACACGTCTG-C3 (IDT) and reverse transcribed with the following indexed primers.

1: NNaaccNNNNNAGATCGGAAGAGCGTCGTGgatcCAGACGTGTGCTCTTC

2: NNacaaNNNNNAGATCGGAAGAGCGTCGTGgatcCAGACGTGTGCTCTTC

3: NNattgNNNNNAGATCGGAAGAGCGTCGTGgatcCAGACGTGTGCTCTTC

4: NNaggtNNNNNAGATCGGAAGAGCGTCGTGgatcCAGACGTGTGCTCTTC

5: NNcgccNNNNNAGATCGGAAGAGCGTCGTGgatcCAGACGTGTGCTCTTC

6: NNccggNNNNNAGATCGGAAGAGCGTCGTGgatcCAGACGTGTGCTCTTC

7: NNctaaNNNNNAGATCGGAAGAGCGTCGTGgatcCAGACGTGTGCTCTTC

8: NNcattNNNNNAGATCGGAAGAGCGTCGTGgatcCAGACGTGTGCTCTTC

9: NNgccaNNNNNAGATCGGAAGAGCGTCGTGgatcCAGACGTGTGCTCTTC

10: NNgaccNNNNNAGATCGGAAGAGCGTCGTGgatcCAGACGTGTGCTCTTC

11: NNggttNNNNNAGATCGGAAGAGCGTCGTGgatcCAGACGTGTGCTCTTC

12: NNgtggNNNNNAGATCGGAAGAGCGTCGTGgatcCAGACGTGTGCTCTTC

13: NNtccgNNNNNAGATCGGAAGAGCGTCGTGgatcCAGACGTGTGCTCTTC

14: NNtgccNNNNNAGATCGGAAGAGCGTCGTGgatcCAGACGTGTGCTCTTC

15: NntattNNNNNAGATCGGAAGAGCGTCGTGgatcCAGACGTGTGCTCTTC

16: NNttaaNNNNNAGATCGGAAGAGCGTCGTGgatcCAGACGTGTGCTCTTC

Resulting cDNAs were circularized and linearized, followed by amplification with the following primers and indexing with unique dual indices (IDT).

Libraries were processed following the workflow in Figure S1 mapping. Briefly, each library was pooled and sequenced on an Illumina HiSeq4000 (Figure 2, Novogene) or Illumina NovaSeq6000 (Figures 1 and 3, UW-Madison Biotechnology Center). Summary statistics and accession numbers are in Supplementary Table 8. Data quality control filtering was done with FastQC v0.11.5, de-multiplexed, adapters were trimmed with BBMAP v37.56, and sequences were aligned using HISAT2 v2.2.1 to hg38/FLUAV for NP eCLIP or a custom hg38/dm6/WSN/VSV genome for spike-in normalized NP and RIG-I eCLIP or Bowtie v2.4.5 to map repetitive elements^79–81^. Trinity was used for RNA-seq *de novo* assembly^82^. PCR duplicates were removed by collapsing reads based on UMIs to yield a unique number of RNAs mapping to each position. The results were used as input for CLIPPER 2.0.1 to identify CLIP peaks statistically enriched over the size-matched input controls^83^. A single size-matched input control was used for analysis of NP CLIP from the doxycycline-induced infected cells and RIG-I CLIP from VSV-infected cells. These samples were pseudoreplicated by randomly splitting reads into two pseudoreplicates for all parts of the workflow that are insensitive to the number of replicates. Data were analyzed with Irreproducible Discovery Rate (IDR) 2.0.03 to measure peak reproducibility across replicates. Bedtools v2.26.0 was used for genomic data manipulation and edgeR was used for comparing across libraries^84,85^. Custom scripts and workflows are available at: https://github.com/mehlelab/.

### Immunofluorescence

tetNP-A549 cells were plated in an 8-well glass bottomed slide (Ibidi) and treated with doxycycline or inoculated with A/WSN/33. Cells were allowed to incubate for 24h, washed in PBS, and crosslinked with 4% paraformaldehyde in PBS. Crosslinked cells were permeabilized with 0.2% Triton-100 in PBS, blocked in 50 mM NH_4_Cl with 4% BSA, and incubated for 1h with 1:1000 dilution anti-RNP (BEI Resources NR-3133). After washing, cells were incubated with 1:1000 donkey anti-goat alexa 568 (ThermoFisher Scientific A11057) and imaged with an EVOS cell imaging system (Thermo Fisher Scientific).

### Cappable-Seq capture of 5’ PPP RNAs

5’ PPP RNAs were selectively labeled with biotinylated cap structure to enable rigorous purification following the Cappable-seq approach^51^. Briefly, RNA was purified from mock or FLUAV-infected A549 cells, reacted with vaccinia capping enzyme (NEB M2080) in the presence of 3’-Desthiobiotin-GTP (NEB N0761), and column purified to fully remove excess 3’-Desthiobiotin-GTP (Zymo Research R1014). RNAs were fragmented by incubation at 95 °C for 2.5 m in the presence of 2.5 mM MgCl_2_. Products were buffer exchanged with AMPure beads (Beckman Coulter A63880) into low TE buffer (10 mM Tris pH 8, 0.1 mM EDTA), captured on hydrophilic streptavidin magnetic beads (NEB S1421S), washed twice in 10 mM Tris pH 7.5, 500 mM NaCl, 1 mM EDTA and twice in 10 mM Tris-HCl pH 7.5, 50 mM NaCl, 1 mM EDTA. RNAs were eluted by incubating for 20 m at room temperature in 10 mM Tris pH 7.5, 50 mM NaCl, 1 mM EDTA and 1 mM biotin. Eluates were buffer exchanged with AMPure beads into low TE. The abundance of specific RNAs in the input and purified fractions was measured by RT-qPCR using primers described above. Fold change in the percent recovery for infected cells versus conditions was calculated.

### Western blotting

Cells were lysed in co-immunoprecipitation buffer supplemented with 0.5% w/v sodium deoxycholate and protease inhibitor cocktail and clarified by centrifugation. Clarified lysates were separated via polyacrylamide gel electrophoresis and transferred to nitrocellulose membranes for blotting. Membranes were blocked and probed with antibodies diluted at the following concentrations: anti-RNP (1:1000, BEI Resources NR-3133), anti-tubulin (1:3000, Sigma T6199), anti-β -actin (1:3000, Proteintech 66009-1-Ig), anti-lamin A/C (Cell Signaling Technologies 2032), or M2 anti-FLAG (1:1000, Sigma A8592). Membranes were imaged on a LiCOR Odyssey Fc with secondary antibodies conjugated to HRP or infrared dyes.

### Quantification and Statistical Analysis

Unless noted in the figure legend, data represent means ± standard deviations (n=3 or more technical replicates) and are representative of at least three independent biological replicates. Boxplots represent the 25^th^-75^th^ percentile and mean and whiskers are calculated by with the Tukey method. Statsmodels, Scipy, Prism and/or matplotlib were used for graphing and statistical analyses. Pairwise comparisons were made using a Student’s T test and multiple comparisons were made using a one-way or two-way ANOVA with the indicated *post hoc* test. Analyses used for CLIP are indicated in their respective methods.

## Notes

### Competing Interest Statement

The authors have declared no competing interest.

### Summary of Updates

Additional experiments with primary human cells reinforce the relevance of the findings. New data are included.

