## Supplementary material for "Influenza virus antagonizes self sensing by RIG-I to enhance viral replication": FIg S

**Figure S1. (Related to Figure 1 and 3). CLIP spike-in controls allow for strict quantitative comparison across experimental conditions of cross-linked RNA.** Cross-linking is performed in human and *Drosophila* cells. The CLIP protocol proceeds as normal up until a size-matched input control is taken. After that, identical amounts of the same *Dropshila* lysate are added as a spike-in control to A549 lysates and the CLIP-seq protocol is completed. This produces sequencing data of both human and *Drosophila* origin. (A) Western blot of RIG-I induction in *Drosophila* cuRIG-I S2 cells and human tetRIG-I A549 cells by addition of $CuCl_2$ for 48 h or 2.5 μg/mL doxycycline for 24h as indicated. (B) Bioinformatic mapping scheme for regular and repetitive sequences followed by read deconvolution for samples with a spike-on control. Note that reads are mapped to a combined dm6/hg38/viral genome and deconvoluted into their respective species after PCR- deduplication. (C) Frequency of reads from spike-in RIG-I CLIP-Seq experiments uniquely mapping to dm6 or hg38 genomes indicate over >99% proper read assignment. (D) Bioinformatic mapping scheme for regular and repetitive sequences during normal CLIP. (E) Western blot of NP induction in *Drosophila* cuNP S2 cells and human tetNP A549 cells by addition of $CuCl_2$ for 48 h or 2.5 μg/mL doxycycline for 24h as indicated. (F) Frequency of reads from spike-in NP CLIP-Seq experiments uniquely mapping to dm6 or hg38 genomes indicate over >99% proper read assignment.

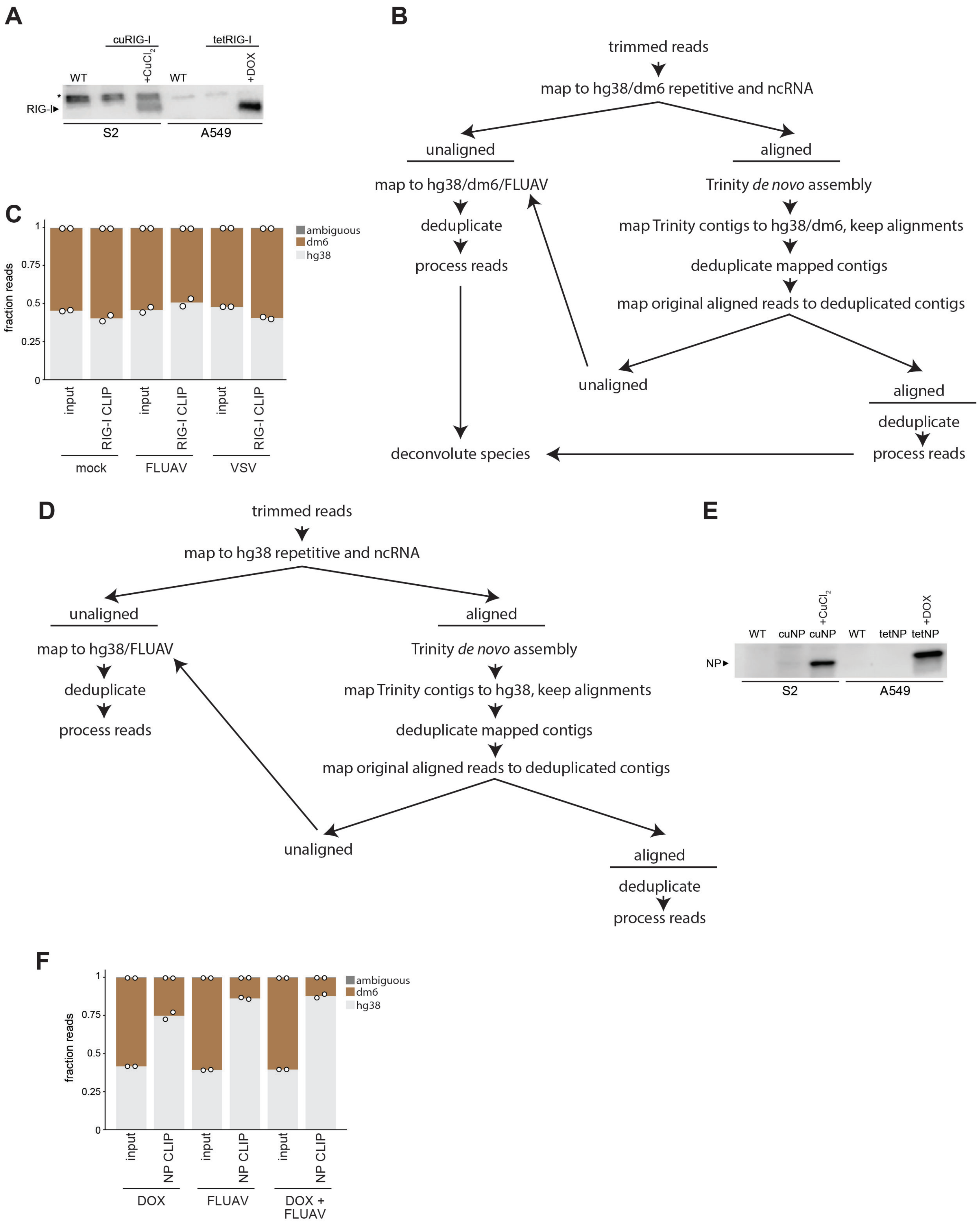
A
cuRIG-I
tetRIG-I
WT
+CuCl2
WT
+DOX
*
RIG-I▸
S2
A549
B
trimmed reads
map to hg38/dm6 repetitive and ncRNA
unaligned
aligned
map to hg38/dm6/FLUAV
Trinity de novo assembly
deduplicate
map Trinity contigs to hg38/dm6, keep alignments
process reads
deduplicate mapped contigs
map original aligned reads to deduplicated contigs
unaligned
aligned
deduplicate
process reads
deconvolute species
C
ambiguous
dm6
hg38
fraction reads
1
0.75
0.5
0.25
0
input
RIG-I CLIP
input
RIG-I CLIP
input
RIG-I CLIP
mock
FLUAV
VSV
D
trimmed reads
map to hg38 repetitive and ncRNA
unaligned
aligned
map to hg38/FLUAV
Trinity de novo assembly
deduplicate
map Trinity contigs to hg38, keep alignments
process reads
deduplicate mapped contigs
map original aligned reads to deduplicated contigs
unaligned
aligned
deduplicate
process reads
E
WT
cuNP
+CuCl2
cuNP
WT
tetNP
+DOX
tetNP
NP▸
S2
A549
F
ambiguous
dm6
hg38
fraction reads
1
0.75
0.5
0.25
0
input
NP CLIP
input
NP CLIP
input
NP CLIP
DOX
FLUAV
DOX +
FLUAV

**Figure S2. (Related to Figure 1). Host transcripts bound by RIG-I remain unchanged between mock and infected conditions.** Spike-in controlled CLIP-Seq was performed for RIG-I in A549 cells stably expressing RIG-I. Cell were in were mock-infected, infected with FLUAV (MOI = 0.01, 24h), or infected with VSV-GFP (MOI = 0.001, 24h). (A) Volcano plot of RIG-I bound transcripts mapped at the gene level showing enrichment of host ncRNAs and tRNAs across all conditions as a function of the false-discovery rate (FDR). (B) Lysates prepared from infected or mock-treated A549 RIG-I cells were used for RIG-I RIPs. The abundance of target RNA was determined by qRT-PCR and shown as fold change in the anti-FLAG (RIG-I) sample versus the IgG control. Data are mean of technical triplicate representative of three biological replicates.

(C) Comparison of RIG-I-bound genes in FLUAV- and mock-infected cells. Log2 fold-change in RIG-I CLIP read depth from FLUAV-infected cells versus mock-infected cells plotted as a function of the average counts per million (CPM) of both libraries.

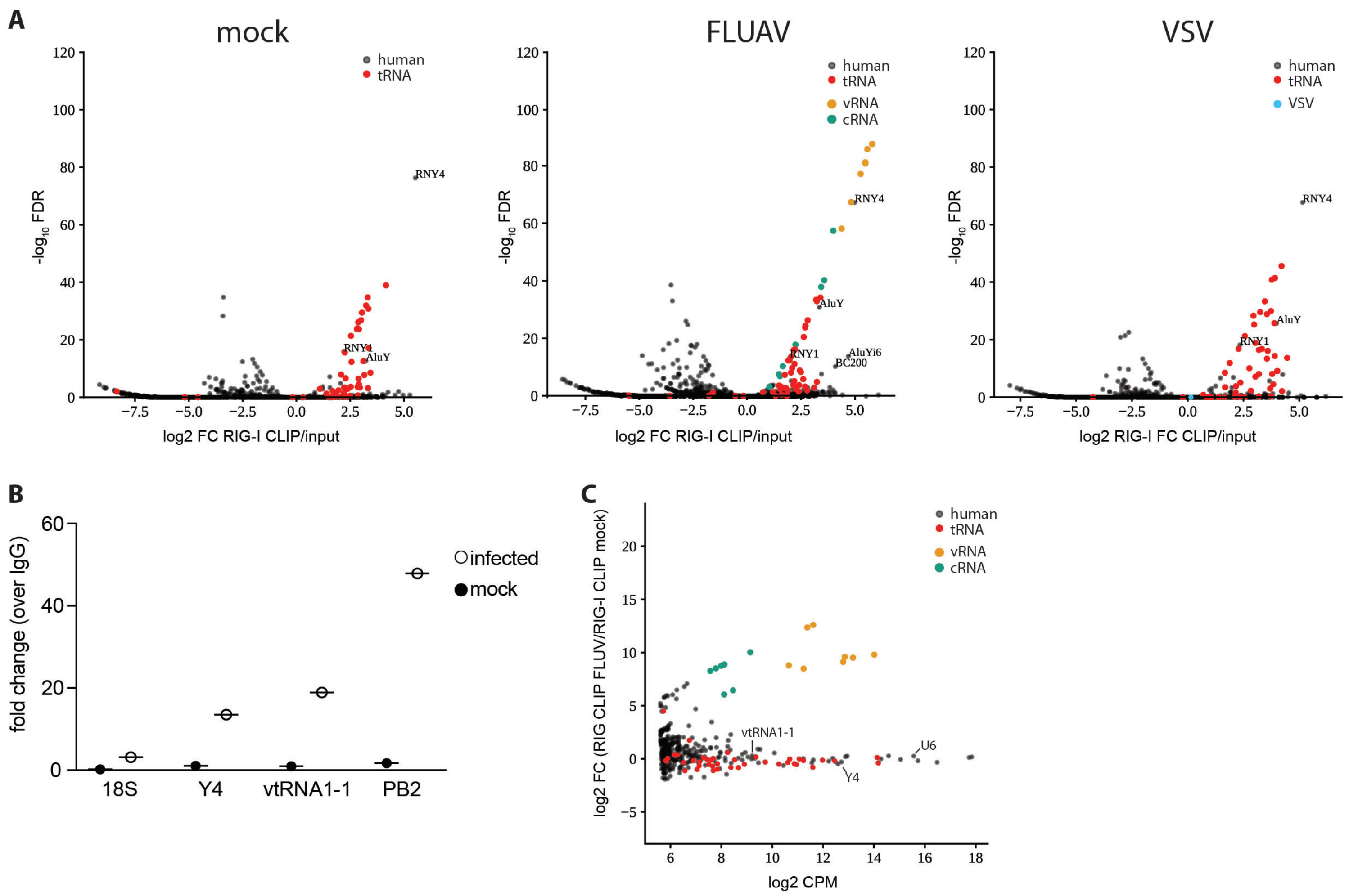

A
mock
FLUAV
VSV
human
tRNA
vRNA
cRNA
VSV
-log10 FDR
log2 FC RIG-I CLIP/input
log2 RIG-I FC CLIP/input
RNY4
RNY1
AluY
AluYi6
BC200
B
fold change (over IgG)
infected
mock
18S
Y4
vtRNA1-1
PB2
C
log2 FC (RIG CLIP FLUV/RIG-I CLIP mock)
log2 CPM
vtRNA1-1
U6
Y4

**Figure S3 (Related to Figure 2). Degradation of RNA in lysates used for co-immunoprecipitation.** RNA was extracted from a fraction of input lysates (left) and immunoprecipitates (right) and RNA integrity was measured with an Agilent Bioanalyzer nano kit.

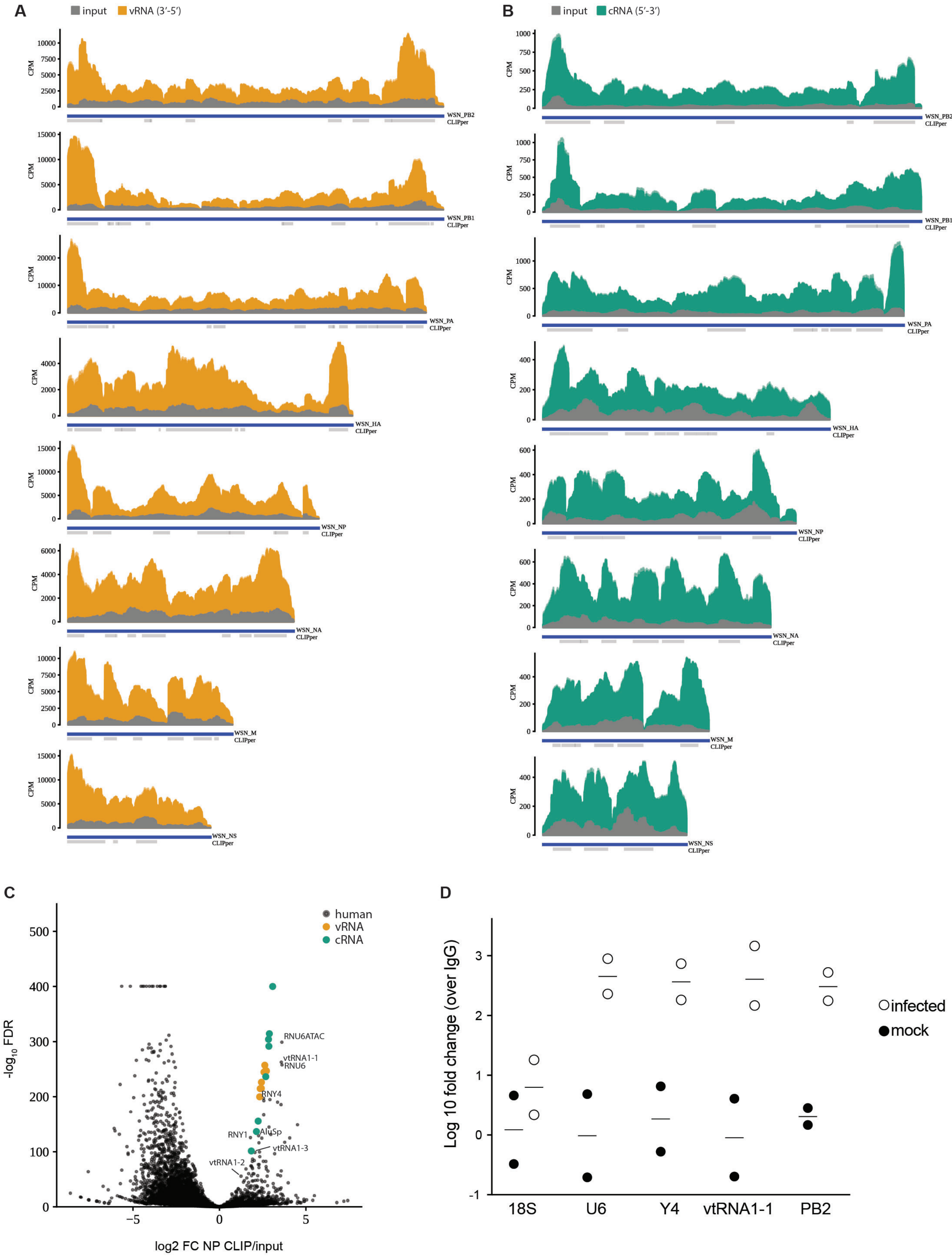
A
input
vRNA (3′-5′)
B
input
cRNA (5′-3′)
CPM
WSN_PB2
WSN_PB1
WSN_PA
WSN_HA
WSN_NP
WSN_NA
WSN_M
WSN_NS
CLIPper
C
human
vRNA
cRNA
RNU6ATAC
vtRNA1-1
RNU6
RNY4
RNY1
AluSp
vtRNA1-3
vtRNA1-2
-log10 FDR
log2 FC NP CLIP/input
D
Log 10 fold change (over IgG)
18S
U6
Y4
vtRNA1-1
PB2
infected
mock

**Figure S4 (Related to Figure 2). NP CLIP recovers crosslinked viral and host RNAs.** (A) Read coverage for NP CLIP libraries in counts per million (CPM) over the negative-sense genomic RNAs (vRNA) or (B) the positive-sense antigenomic RNAs (cRNAs) demonstrating uneven coverage across genome. Data from independent biological replicates are shown as mean and s.d. using dark and light shades of the same color, respectively. CLIPper-called peaks are denoted along the bottom of the read track. (C) NP binds host RNAs. Volcano plot of NP-bound RNAs mapped at the gene level showing enrichment of viral genomes and host ncRNAs as a function of the false-discovery rate (FDR). Associated with Supplementary Table 4. (D) Lysates prepared from infected or mock-treated A549 cells were used for NP RIPs. The abundance of target RNA was determined by qRT-PCR and shown as fold change in the anti-NP sample versus the IgG control. Data are mean from two independent biological replicates performed in triplicate.

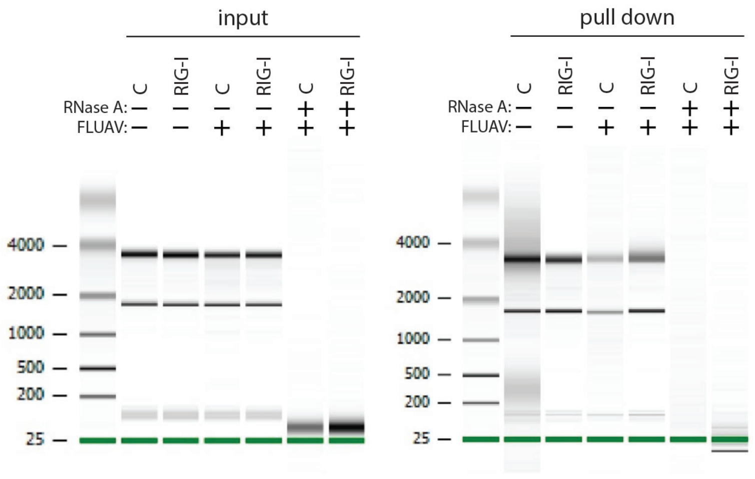
input
pull down
C
RIG-I
RNase A:
FLUAV:
4000
2000
1000
500
200
25

**Figure S5. (Related to Figure 3) Host ncRNA are bound by NP in the cytoplasm** (A) NP CLIP-seq with a spike-in control was performed in A549 cells where NP was expressed via doxycycline induction or infection with FLUAV (MOI = 0.01. 24 h). Volcano plot of RNAs bound mapped at the gene level. The zoom-in highlights host ncRNAs bound by NP during infection. (B) Cytoplasmic and nuclear extracts were prepared from infected A549 cells. Fraction purity was monitored by blotting for tublin, lamin, and NP. (C) Host ncRNAs bound by NP during infection mirror those bound by cytoplasmic NP. NP CLIP-seq was performed on cytoplasmic and nuclear fractions and log2 fold-change in read density at the gene level was calculated between conditions. Similar analysis was performed for the data presented in (A). Both comparisons were plotted against each other with select host ncRNAs highlighted. (D) Principal component analysis of NP-bound RNAs in whole cell lysates or subcellular fractions. Samples were integrated from two different experiments (Batch 1 and 2). PCo1 separates samples by observed or biochemically fractionated localization of NP. (E) UpSet plot showing shared RNA-binding peaks between NP and RIG-I analysis restricted to host ncRNAs and viral RNAs under different experimental conditions.

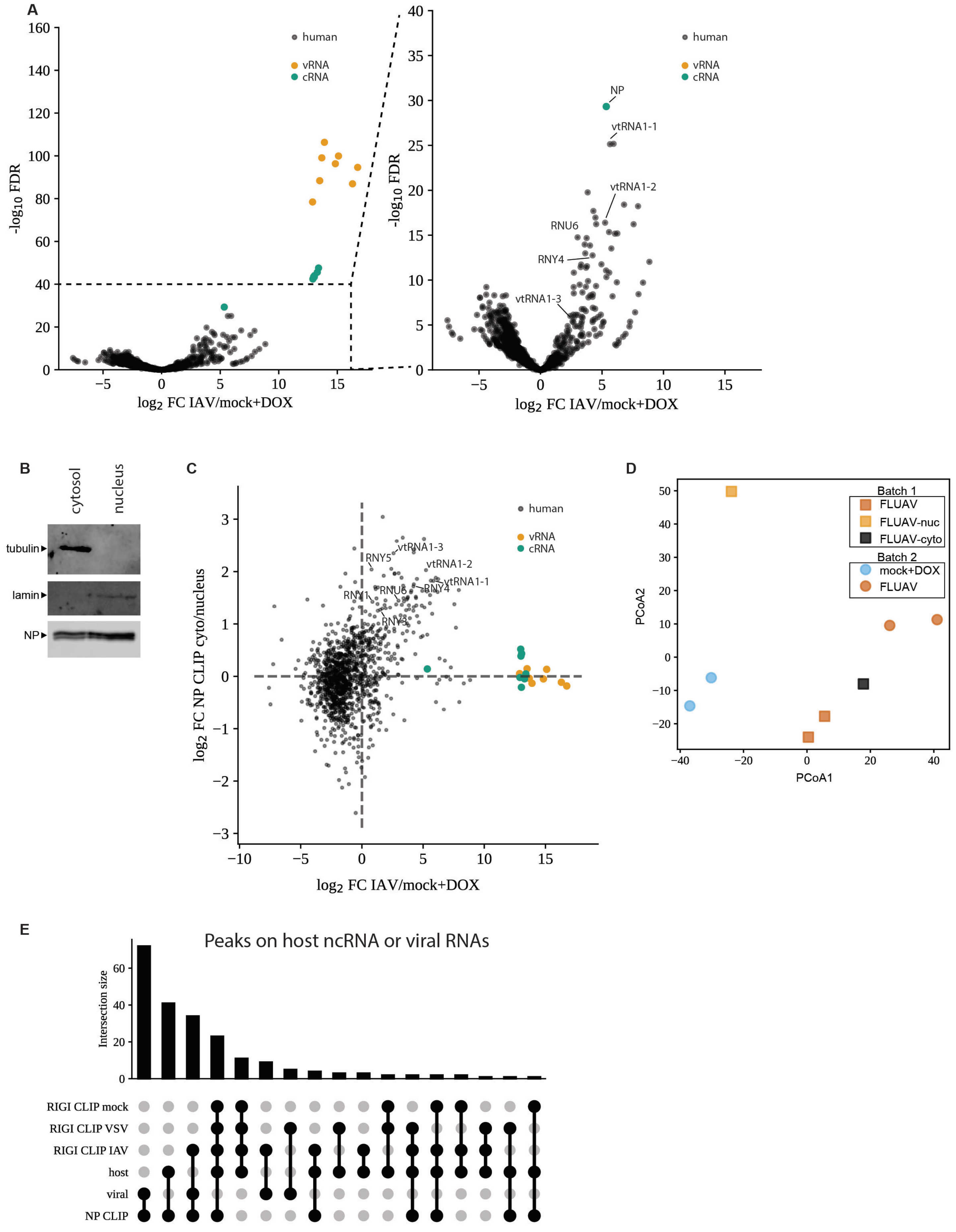
A
human
vRNA
cRNA
-log10 FDR
log2 FC IAV/mock+DOX
NP
vtRNA1-1
vtRNA1-2
RNU6
RNY4
vtRNA1-3
B
cytosol
nucleus
tubulin
lamin
NP
C
log2 FC NP CLIP cyto/nucleus
RNY5
vtRNA1-3
vtRNA1-2
vtRNA1-1
RNY1
RNU6
RNY4
RNY3
D
Batch 1
FLUAV
FLUAV-nuc
FLUAV-cyto
Batch 2
mock+DOX
FLUAV
PCoA2
PCoA1
E
Peaks on host ncRNA or viral RNAs
Intersection size
RIGI CLIP mock
RIGI CLIP VSV
RIGI CLIP IAV
host
viral
NP CLIP

**Figure S6 (Related to Figure 4-5) RNase-H depletion of immunostimulatory influenza genomic RNAs and NP localization.** (A) Total eukaryotic RNA or *in vitro* transcribed influenza genomic vRNAs were treated with an RNase-H-based depletion protocol targeting influenza genomic RNAs or mock treated. Resultant RNAs were transfected into ISRE-reporter cells and assayed for ISRE stimulation. ** $P < 0.01$ by one-way ANOVA with Dunnett's multiple comparisons test against mock-transfected cells. (B) Eluates of NP RNA-immunoprecipitations from FLUAV infected A549s (MOI = 0.02, 24h), IgG controls, or *in vitro* transcribed influenza genomic RNAs were treated with the RNase-H based depletion protocol. RNA were separated via TBE urea polyacrylamide gel electrophoresis and imaged with SYBR-gold staining. (C) WT, RIG knockout (*DDX58*$^{-/-}$), or MAVS knockout (*MAVS*$^{-/-}$) A549 IFN-β-reporter cells were transfected with decreasing amounts of *in vitro* transcribed non-coding RNAs, poly IC, or mock transfected. IFN-β promoter induction shown relative to mock-treated conditions for each cell line. *** $P < 0.001$ as determined by a two-way ANOVA with *post hoc* Tukey's multiple comparisons test. (D) Subcellular localization of WT or mutant NP-GFP fusion proteins were visualized by fluorescence microscopy.

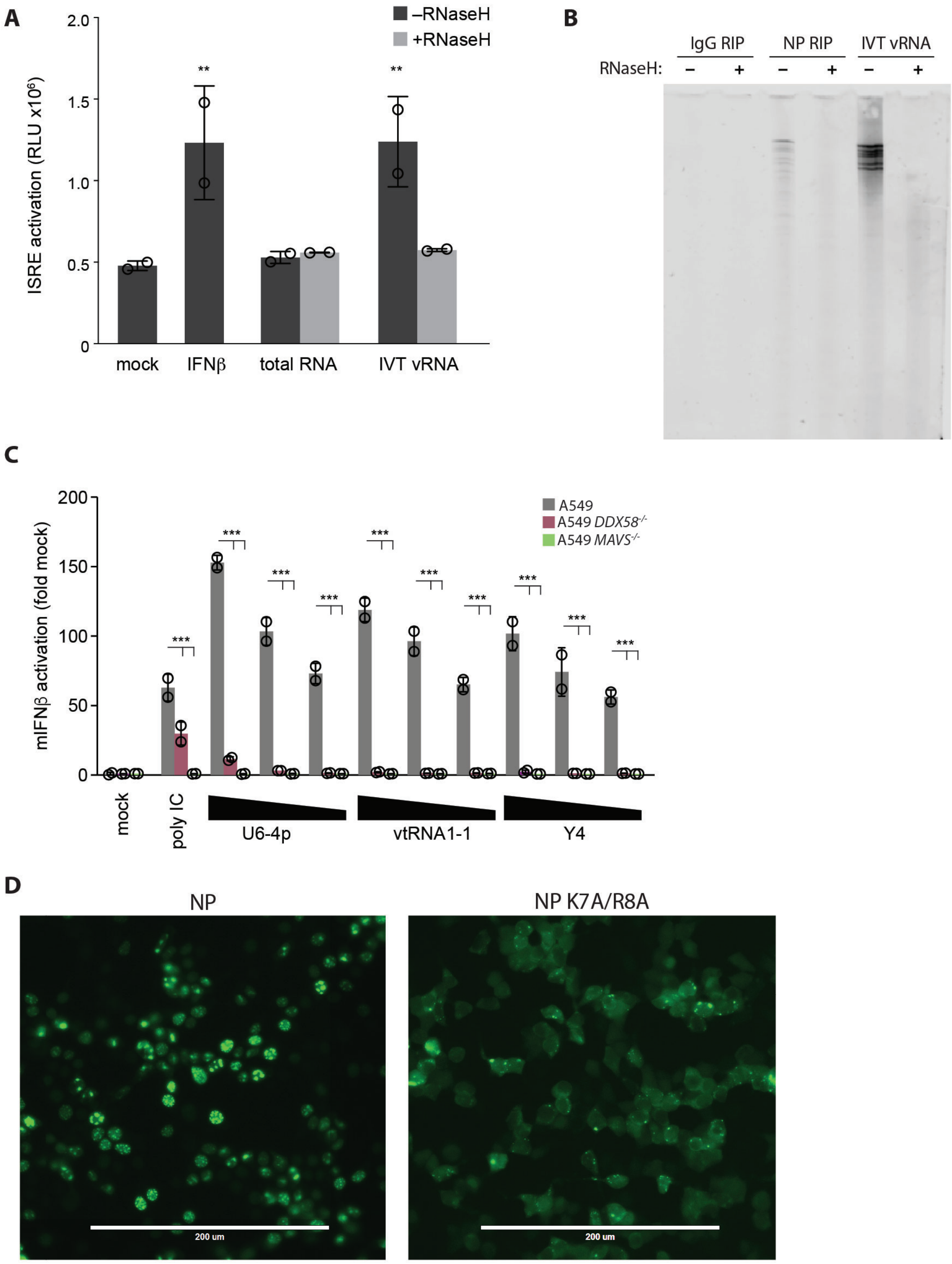
A
–RNaseH
+RNaseH
ISRE activation (RLU x10^6)
2.0
1.5
1.0
0.5
0
**
**
mock
IFNβ
total RNA
IVT vRNA
B
IgG RIP
NP RIP
IVT vRNA
RNaseH:
–
+
–
+
–
+
C
A549
A549 DDX58-/-
A549 MAVS-/-
mIFNβ activation (fold mock)
200
150
100
50
0
***
mock
poly IC
U6-4p
vtRNA1-1
Y4
D
NP
NP K7A/R8A
200 um
200 um

**Figure S7(Related to Figure 5) Non-coding RNAs purified from primary human cells infected with the pandemic isolate A/California/04/2009 activate innate immunity.** (A) RNA was extracted from normal human bronchial epithelial (NHBE) cells infected with a primary influenza A isolates (A/California/04/2009 MOI = 0.5). RNA was used for ASO purifications, purified RNAs were reintroduced into A549 ISRE-NLuc2AGFP reporter cells, and cells were imaged for GFP fluorescence 24 h later. These cells were then infected and viral titers are reported in Fig 5H. (B) Normal human bronchial epithelial (NHBE) cells differentiated at the air-liquid interface (ALI) were infected with A/California/04/2009 or mock treated. Infected cells were isolated by flow cytometry prior to extraction of total RNA. Target RNAs were purified with ASO, reintroduced into A549 ISRE-NLuc2AGFP reporter cells, and cells were imaged for GFP fluorescence 24 h later. These cells were then infected and viral titers are reported in Fig 5I. Images for each technical replicate are shown. Scale bar = 400 $\mu$m.

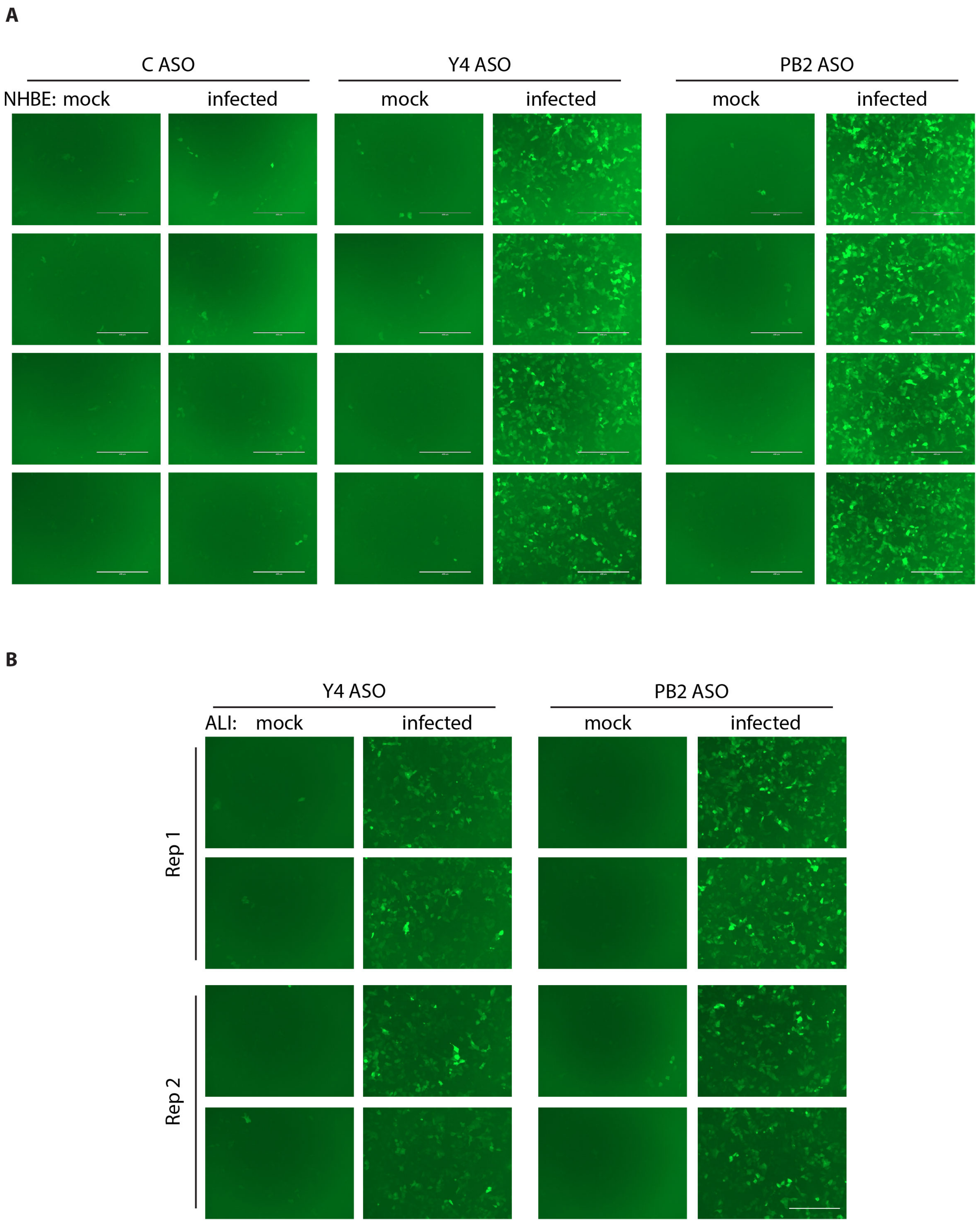
A
C ASO
Y4 ASO
PB2 ASO
NHBE: mock
infected
mock
infected
mock
infected
B
Y4 ASO
PB2 ASO
ALI: mock
infected
mock
infected
Rep 1
Rep 2

**Figure S8 Uncropped blots**

Fig 1A

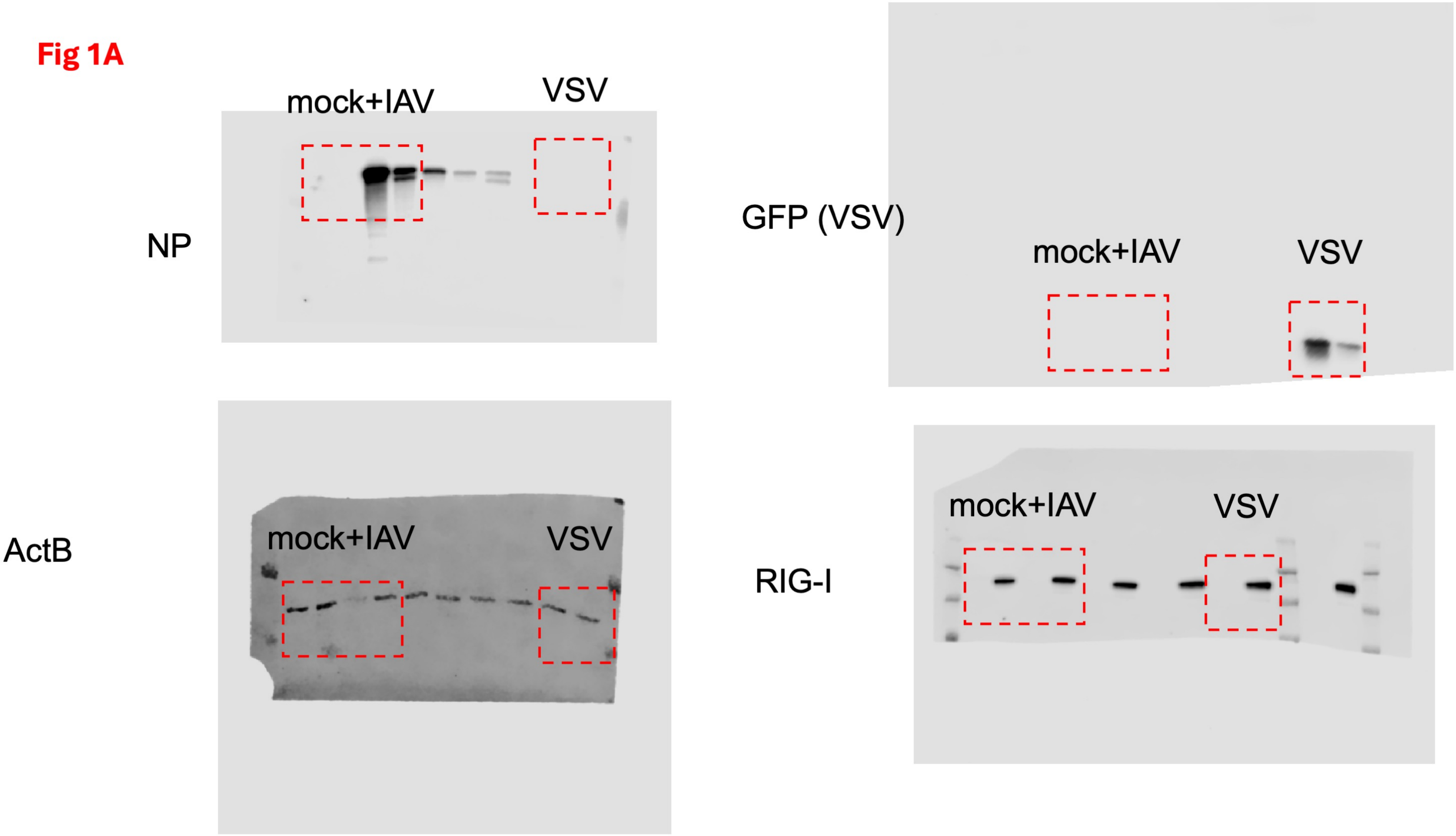

mock+IAV
VSV
NP
GFP (VSV)
mock+IAV
VSV
ActB
mock+IAV
VSV
RIG-I
mock+IAV
VSV

**2A**

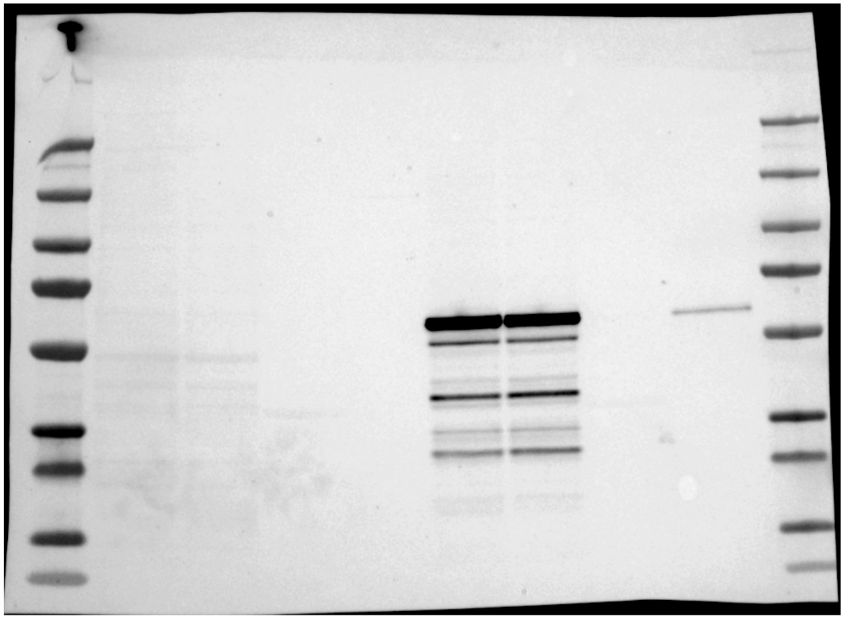

anti-NP

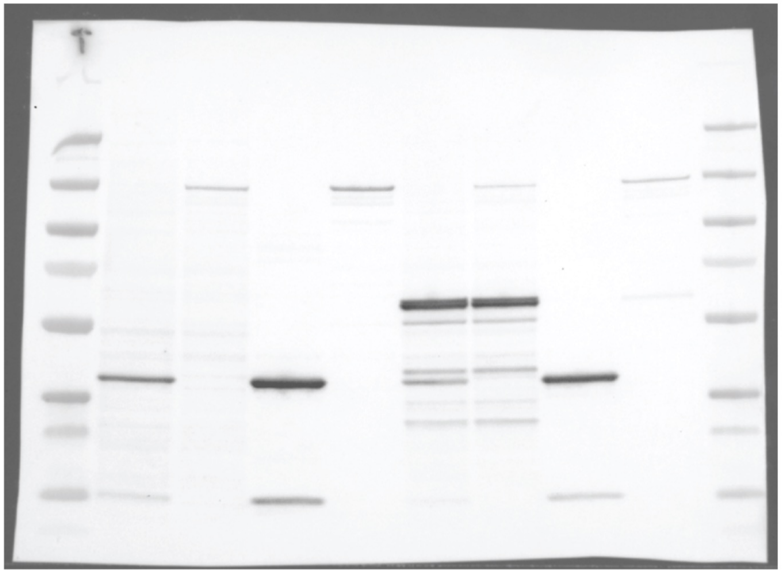

anti-Strep
anti-NP

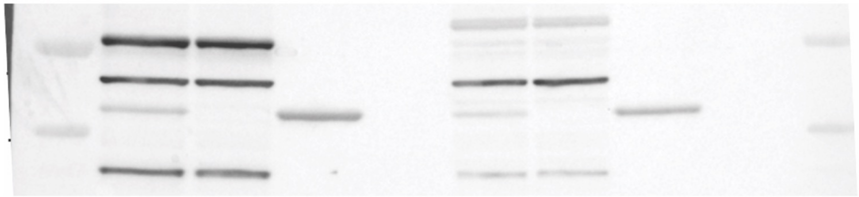

anti-actin

**2B**

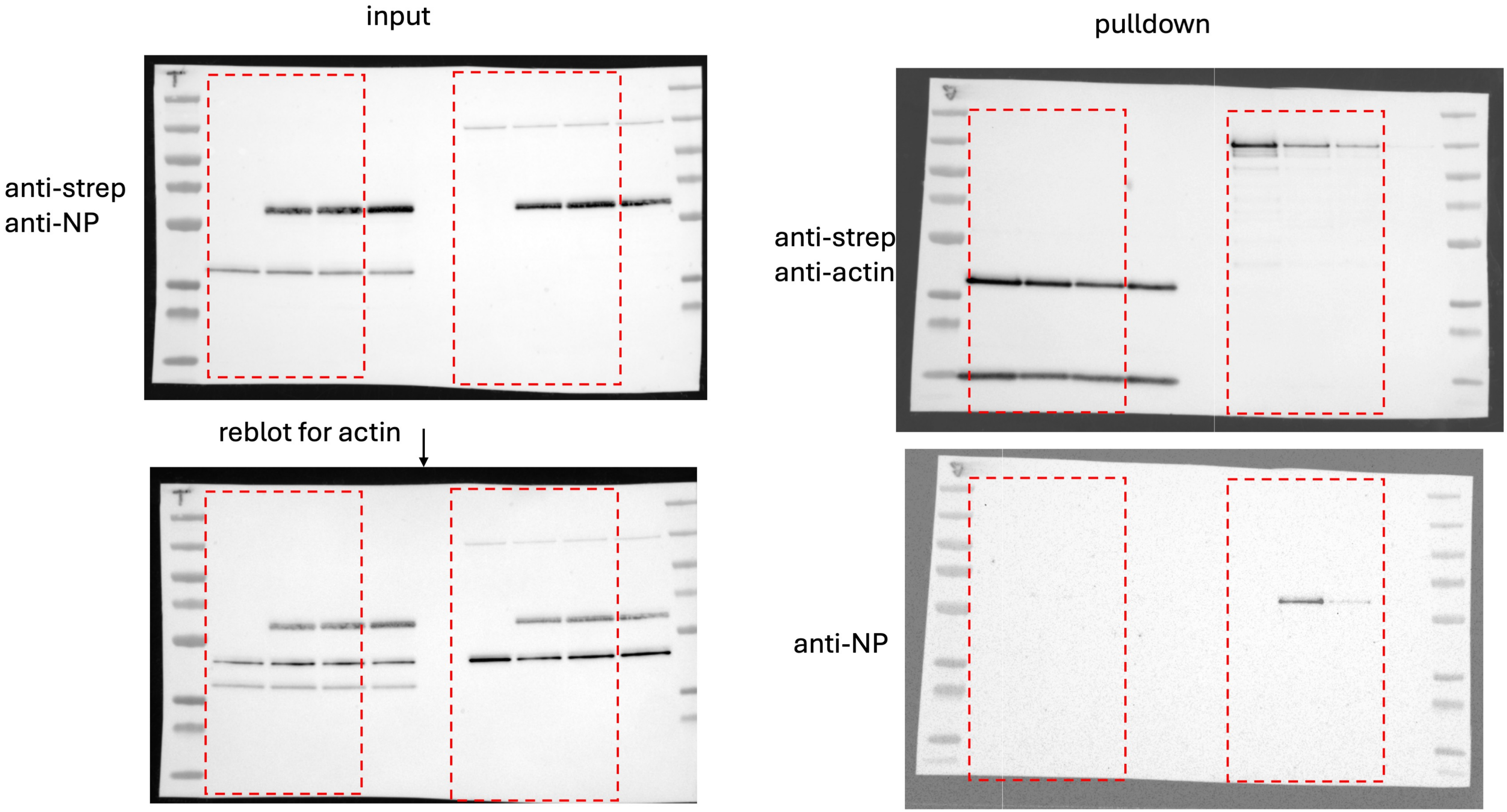
input
pulldown
anti-strep
anti-NP
reblot for actin
anti-strep
anti-actin
anti-NP

2C

NP CLIP

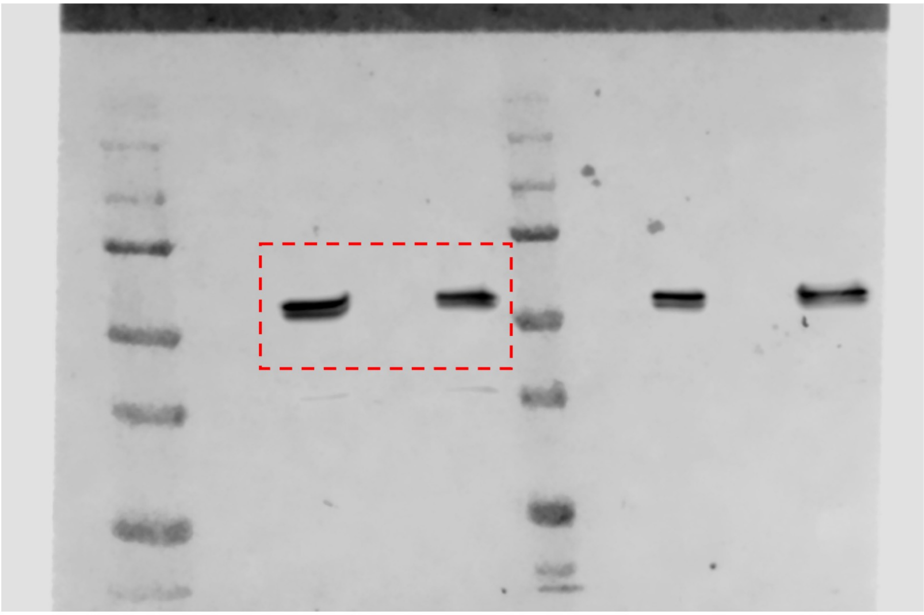

Fig 3A

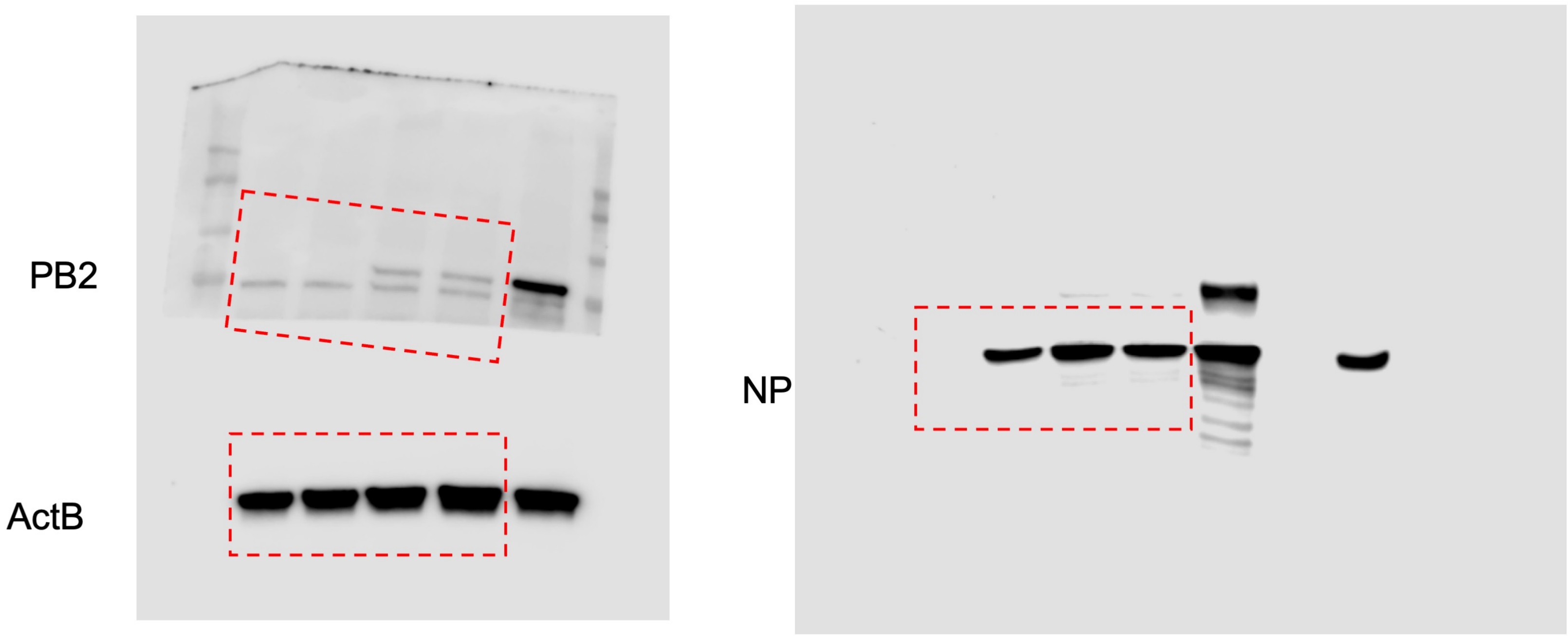

**Fig 3G**

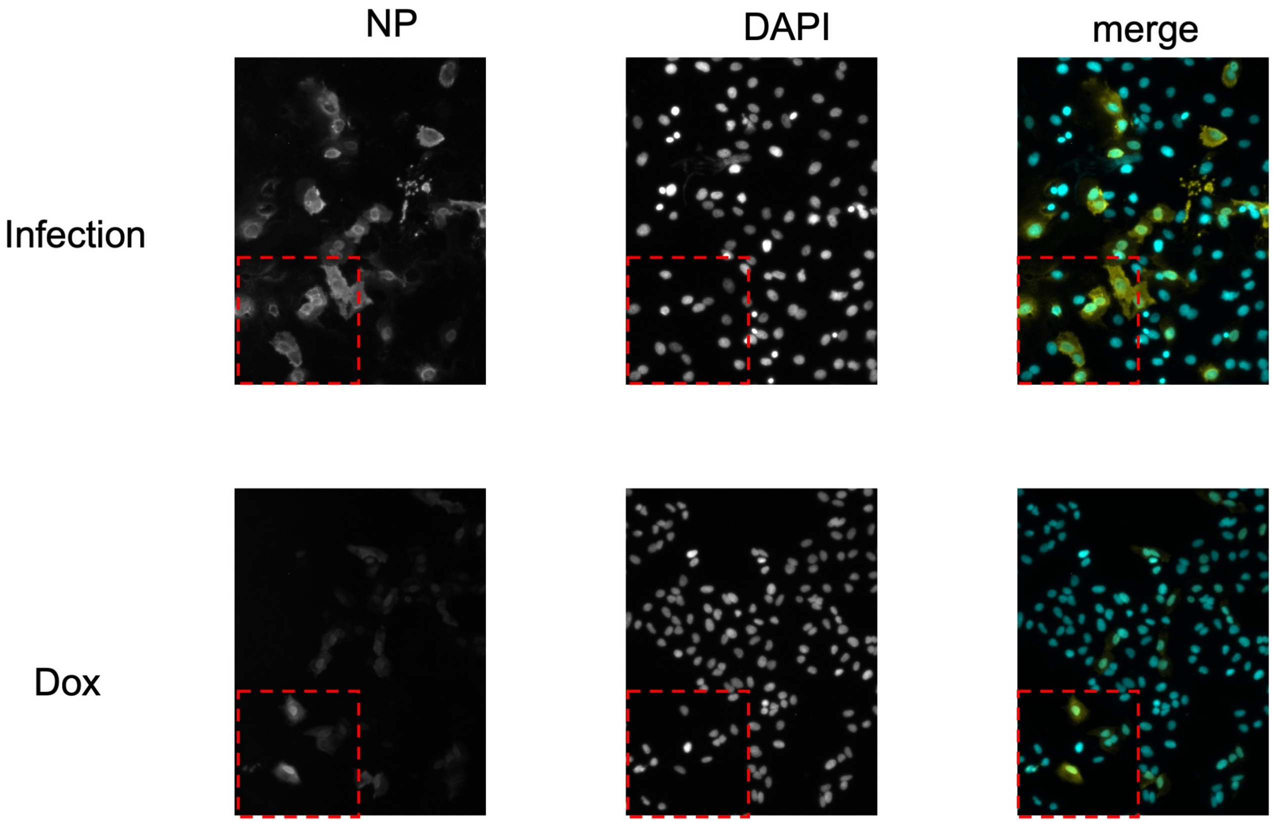
